# Overlooked tertiary sulci serve as a meso-scale link between microstructural and functional properties of human lateral prefrontal cortex

**DOI:** 10.1101/2020.03.24.006577

**Authors:** Jacob A. Miller, Willa I. Voorhies, Daniel J. Lurie, Mark D’Esposito, Kevin S. Weiner

## Abstract

Understanding the relationship between neuroanatomy and function in portions of human cortex that are expanded compared to other mammals such as lateral prefrontal cortex (LPFC) is of major interest in systems and cognitive neuroscience. When considering neuroanatomical-functional relationships in LPFC, shallow indentations in cortex known as tertiary sulci have been largely ignored. Here, by implementing a multi-modal approach and manually defining 936 neuroanatomical structures in 72 hemispheres (males and females), we show that a subset of these overlooked tertiary sulci serve as a meso-scale link between microstructural (myelin content) and functional (network connectivity) properties of human LPFC in individuals. For example, the *posterior middle frontal sulcus* (*pmfs*) is a tertiary sulcus with three components that differ in their myelin content, resting state connectivity profiles, and engagement across meta-analyses of 83 cognitive tasks. Further, generating microstructural profiles of myelin content across cortical depths for each *pmfs* component and the surrounding middle frontal gyrus (MFG) shows that both gyral and sulcal components of the MFG have greater myelin content in deeper compared to superficial layers and that the myelin content in superficial layers of the gyral components is greater than sulcal components. These findings support a classic, yet largely unconsidered theory that tertiary sulci may serve as landmarks in association cortices, as well as a modern cognitive neuroscience theory proposing a functional hierarchy in LPFC. As there is a growing need for computational tools that automatically define tertiary sulci throughout cortex, we share *pmfs* probabilistic sulcal maps with the field.

**Significance statement:** Lateral prefrontal cortex (LPFC) is critical for higher-order cognitive control and goal-directed behavior and is disproportionately expanded in the human brain. However, relationships between fine-scale neuroanatomical structures largely specific to hominoid cortex and functional properties of LPFC remain elusive. Here, we show that these structures, which have been largely neglected throughout history, surprisingly serve as markers for anatomical and functional organization in human LPFC. These findings have theoretical, methodological, developmental, and evolutionary implications for improved understanding of neuroanatomical-functional relationships not only in LPFC, but also in association cortices more broadly. Finally, these findings ignite new questions regarding how morphological features of these neglected neuroanatomical structures contribute to functions of association cortices that are critical for human-specific aspects of cognition.

## Introduction

Understanding how anatomical structures of the brain support functional gradients and networks that perform computations for human-specific aspects of cognition is a major goal in systems and cognitive neuroscience. Of the many anatomical structures to target, lateral prefrontal cortex (LPFC), which is disproportionately expanded in the human brain, is particularly important given its central role in cognitive control and goal-directed behavior (Miller and Cohen, 2001; Szczepanski and Knight, 2014; Donahue et al., 2018). Major progress has been made in understanding the relationship between the functional organization and the large-scale cortical anatomy of human LPFC. For example, modern neuroimaging research shows widespread support for a hierarchical functional gradient organized along the rostral-caudal anatomical dimension of LPFC spanning several centimeters (Badre and D’Esposito, 2009; Nee and D’Esposito, 2016; Demirtas et al., 2019). Beyond this large-scale organization of human LPFC, it is unknown if more fine-grained structural-functional relationships exist. Thus, to begin to fill this gap in knowledge, we sought to answer the following question in the present study: Do individual differences in fine-grained morphological features of LPFC shed light on microstructural and functional properties of LPFC?

An important morphological feature of cortex is the patterning of the indentations, or sulci. Indeed, 60-70% of the cortex is buried in sulci and some sulci serve as landmarks that identify different cortical areas, especially in primary sensory cortices (Van Essen and Dierker, 2007; Zilles et al., 2013). In these cases, merely identifying a sulcus provides functional insight (Hinds et al., 2008). Despite this widely replicated relationship between sulcal morphology and functional representations in primary sensory cortices, much less is known regarding the predictability between shallow, tertiary sulci and functional representations in association cortex, especially LPFC. A classic theory proposed by Sanides (1964) hypothesized that the late emergence and protracted development of tertiary sulci may co-occur with microstructural and functional features of association cortices, as well as cognitive functions such as working memory or inhibitory control, that each also show a protracted development (Curtis and D’Esposito, 2003; Petrides, 2005; Gazzaley and Nobre, 2012).

However, at least two factors have prevented the examination of tertiary sulci relative to anatomical and functional organization in human LPFC. First, tertiary sulci are presently excluded from nearly all published neuroanatomical atlases because classic anatomists could not discriminate tertiary sulci from indentations produced by veins and arteries on the outer surface of the cerebrum in post-mortem tissue, which is considered the gold standard of anatomical research (Weiner et al., 2018). Consequently, the patterning of tertiary sulci within LPFC has a contentious history, whereby sulci in the posterior middle frontal gyrus (MFG) were either undefined in classic atlases or conflated with more anterior structures (**Figure 1**). Second, the majority of human functional magnetic resonance imaging (MRI) studies implement group analyses on average brain templates. As shown in **Figure 1**, averaging cortical surfaces together causes tertiary sulci in LPFC to disappear, especially within the posterior MFG.

**Figure 1.**
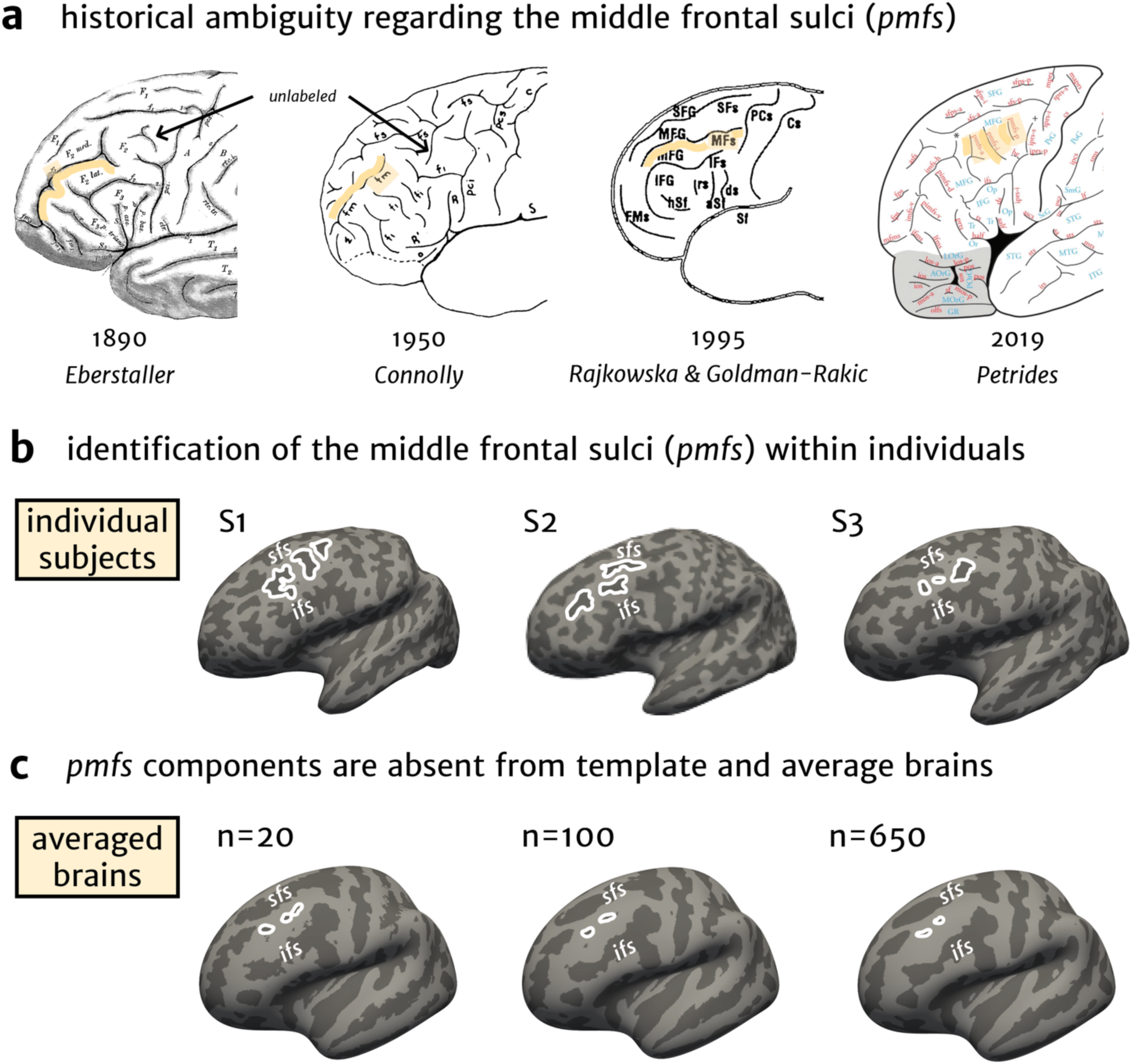
A synopsis of ambiguity regarding sulcal definitions in the human posterior middle frontal gyrus over the last 130 years. Classic and modern schematics of the sulcal patterning in human lateral prefrontal cortex (LPFC). (**a**) Sulci in the middle frontal gyrus are labeled in yellow on classic and modern schematics of human LPFC. Historically, anatomists had previously either (1) not labeled the sulci within the location of the modern *pmfs* (first two images; arrow indicates depicted, but unlabeled sulcal components) (Eberstaller, 1890; Connolly, 1950) or 2) included these sulci in the definition of the posterior portion of the frontomarginal sulcus (third image; (Rajkowska and Goldman-Rakic, 1995). The most recent schematic (fourth image) proposes that the *pmfs* is separate from the intermediate frontal sulcus (*imfs-h and imfs-v*, synonymous with the *frontomarginal sulcus*) and consists of three distinct components: posterior (*pmfs-p*), intermediate (*pmfs-i*), and anterior (*pmfs-a*). (**b**) Three individually labeled left hemispheres with the *pmfs* outlined in white. The *pmfs* is prominent within individual subjects (**Extended Data Fig. 2-1** for all subjects). The superior and inferior frontal sulci (*sfs, ifs*) are labeled for reference above and below the middle frontal gyrus, respectively. (**c**) Average cortical surfaces show much smaller *pmfs* components compared to individual subjects. As more subjects are averaged together into templates, the *pmfs* disappears almost entirely, which is inconsistent with their prominence in individual hemispheres.

Here, we implemented a multi-modal approach demonstrating that carefully identifying individual sulci in LPFC reveals that the *posterior middle frontal sulcus* (*pmfs*) serves as a meso-scale link between microstructural (myelin content) and functional (network connectivity) properties of human LPFC in individual participants. We applied a recently proposed labeling scheme of tertiary sulci in LPFC (Petrides and Pandya, 2012; Petrides, 2019) from post-mortem brains to test whether these sulci could be defined in the LPFC of individual subjects *in-vivo* using MRI. We find that three components of the *pmfs* are dissociable based on myelin content, resting state functional connectivity profiles, and cognitive task activations. Moreover, the *pmfs* shows a distinct microstructural profile of myelin content across cortical depths from the surrounding MFG. Together, these results not only provide important evidence that individual differences in LPFC sulcal patterning reflects meaningful differences in microstructural and functional properties, but also suggest the *pmfs* serves as a bridge to Sanides’ classic hypothesis.

## Materials and Methods

In the sections below, we describe the data used and the analysis methods implemented in three separate sections: 1) the general approach and a description of the multi-modal datasets that were used, 2) a detailed description of the methodology used for sulcal labeling within individual subjects, and 3) the calculation of anatomical and functional metrics.

### General approach

We sought to characterize sulcal morphology at the individual subject level in the LPFC of the human brain. To implement this process, we manually defined sulci following the most recent and comprehensive proposed labeling of sulci in the frontal lobe based on post-mortem specimens (Petrides and Pandya, 2012; Petrides, 2019). As in our prior work (Weiner et al., 2014), all sulci were defined in native space cortical surfaces and individual hemispheres, which enables the most accurate definition of tertiary sulci in *in-vivo* MRI data.

### Multi-modal HCP dataset

We analyzed a subset of the multi-modal MRI data available for individual subjects from the Human Connectome Project (HCP). We began with the first 5 numerically listed HCP subjects and then randomly selected 31 additional human subjects from the HCP for a total of 36 individuals (17 female, 19 male, age range 22-36 years).

Anatomical T_1_-weighted (T_1_w) MRI scans (0.8 mm voxel resolution) were obtained in native space from the HCP database, along with outputs from the FreeSurfer pipeline slightly modified by the HCP (Dale et al., 1999; Fischl et al., 1999a; Glasser et al., 2013). Maps of the ratio of T_1_-weighted and T_2_-weighted scans, which is a measure of tissue contrast enhancement related to myelin content, were downloaded as part of the HCP ‘Structural Extended’ release. All additional anatomical metrics, which are detailed in the next section, were calculated on the full-resolution, native FreeSurfer (https://surfer.nmr.mgh.harvard.edu/) meshes (Dale et al., 1999; Fischl et al., 1999a; Fischl et al., 1999b).

#### Anatomical labeling and metrics

### Manual sulcal labeling

Guided by a recent comprehensive proposal for labeling sulci in LPFC, each sulcus was manually defined within each individual hemisphere on the FreeSurfer *inflated* mesh with *tksurfer*. The *curvature* metric in FreeSurfer distinguished the boundaries between sulcal and gyral components, and manual lines were drawn to separate sulcal components based upon the proposal by Petrides and colleagues (2012; 2019), as well as the appearance of sulci across the *inflated, pial*, and *smoothwm* surfaces. We maintained the number of components for all sulci (e.g., the three components of the *posterior* m*iddle frontal sulcus - pmfs*) based on the proposal by Petrides and colleagues to test if each of these sulcal components could be defined in a relatively large sample size (N=72) of *in-vivo* hemispheres. That is, the proposal for defining sulci in LPFC that is included in the book chapter (Petrides and Pandya, 2012) and most recent atlas (Petrides, 2019) are summarized as a schematic. It is unclear from which data (or series of data) this schematic was generated as - to our knowledge - there have been no empirical papers published with these sulcal definitions and morphological analyses in individual hemispheres until the present work. Consequently, our definitions were identical to the proposal in order to test whether this distinction among sulci that is summarized in the schematic could be applied to empirical data in individual hemispheres. The labels were generated using a two-tiered procedure. The labels were first defined manually by J.M. and W.V. and then finalized by a neuroanatomist (K.S.W.). All anatomical labels for a given hemisphere were fully defined before any morphological or functional analysis of the sulcal labels was performed. The superior, inferior, posterior, and anterior boundaries of our cortical expanse of interest were the following sulci, respectively: (1) the anterior and posterior components of the *superior frontal sulcus*, (2) the *inferior frontal sulcus*, (3) the *central sulcus*, and (4) the horizontal (*imfs-h*) and vertical (*imfs-v*) intermediate frontal sulci. An example hemisphere with every sulcus labeled within these boundaries is shown in **Figure 2a**, and the *pmfs* sulcal components are plotted on each hemisphere in **Extended Data Fig. 2-1**.

**Figure 2.**
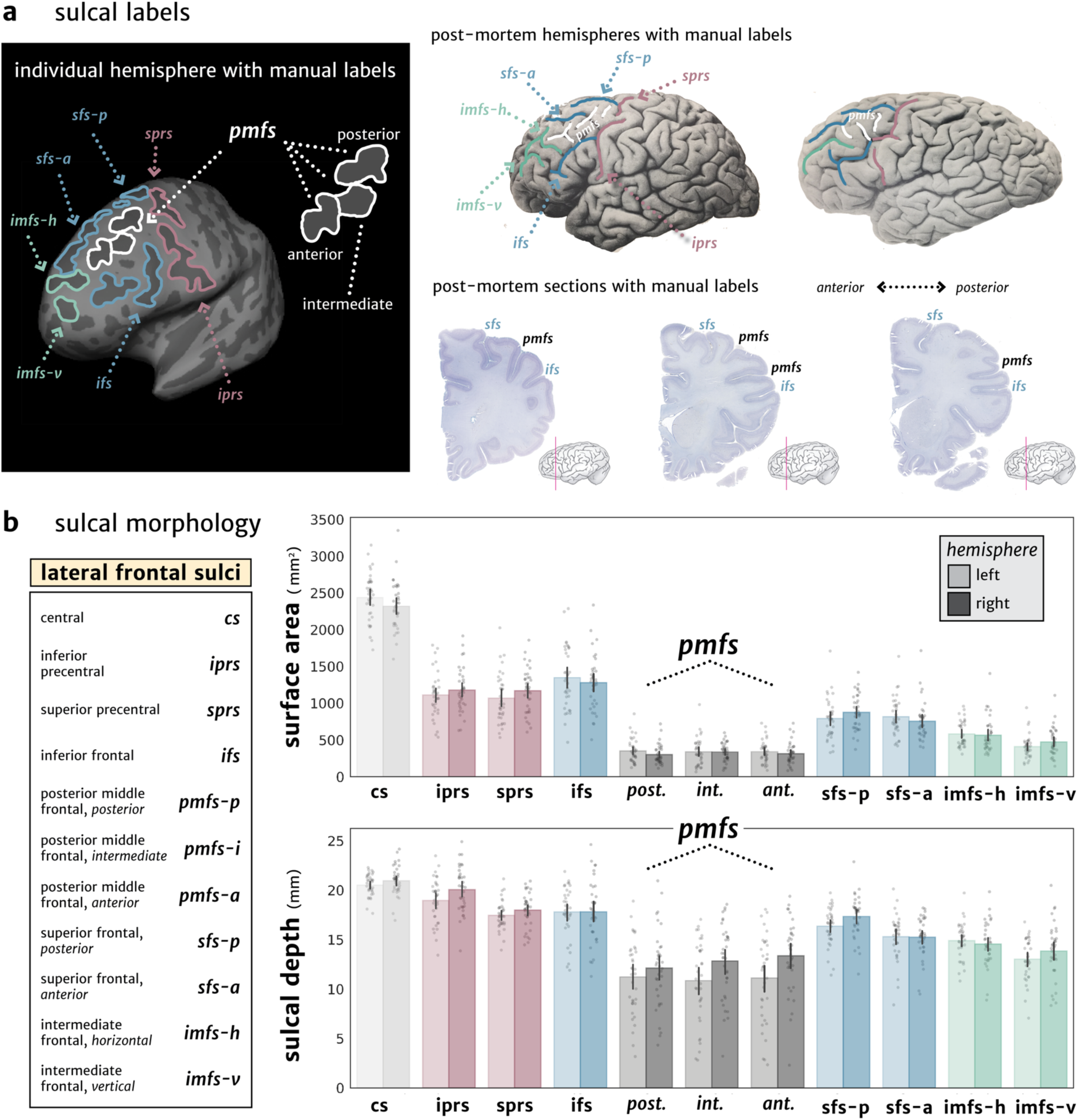
LPFC tertiary sulci are easily identifiable and characteristically shallow. (**a**) Left, an example inflated cortical surface of an individual left hemisphere in which the sulci examined in the present study are outlined and labeled. Sulci are dark gray, while gyri are light gray. Right, two different post-mortem brains (Retzius, 1896) and three histological sections (note that the pmfs components are referred to as “intermediate frontal sulcus” in the Allen Human Brain Atlas: https://atlas.brain-map.org/, (Ding et al., 2016)) showing that the *pmfs* sulci are also identifiable in post-mortem tissue samples. (**b**) Top: Surface area for each sulcus (ordered posterior to anterior) is plotted for each individual subject (gray circles), as well as the mean (colored bars) and 95% confidence interval (black line). Acronyms used for each LPFC sulcus are also included. Darker shades indicate right hemisphere values, while lighter shades indicate left hemisphere values. The three *pmfs* sulci have the smallest surface area of all sulci in LPFC. Bottom: Same layout as above, but for sulcal depth (mm), as calculated from a recent algorithm (Madan, 2019). The three *pmfs* sulci are the shallowest of the LPFC sulci measured here.

### Quantification of sulcal depth and surface area

Sulcal depth was calculated from the native meshes generated by the FreeSurfer HCP pipeline. Raw values for sulcal depth (mm) were calculated from the sulcal fundus to the smoothed outer pial surface using a custom-modified version of a recently developed algorithm for robust morphological statistics building on the FreeSurfer pipeline (Madan, 2019). Surface area (mm^2^) was generated for each sulcus through the *mris_anatomical_stats* function in FreeSurfer (Dale et al., 1999; Fischl et al., 1999a). We focused on sulcal depth as it is the main measurement that is used to discriminate tertiary sulci from primary and secondary sulci. Specifically, primary sulci are deepest, while tertiary sulci are shallowest, and secondary sulci are in between (Welker, 1990). We also included surface area as tertiary sulci typically also have a reduced surface area compared to primary and secondary sulci.

### Calculating T_1_w/T_2_w myelin index along an anterior-posterior dimension in LPFC

In order to test if there is a relationship between any of our sulci of interest and myelination content, we used an *in vivo* proxy of myelination: the T_1_w/T_2_w maps for each individual hemisphere (Glasser and Van Essen, 2011; Shams et al., 2019). We averaged this value across each vertex for each sulcus in order to demonstrate if the *pmfs* sulcal components are separable based on myelin content (**Figure 3**). We further sought to characterize the relationship between morphology and myelin by determining if there was an anterior-posterior gradient of myelination across individual hemispheres. First, for each individual hemisphere, we calculated the minimum geodesic distance of each vertex from the central sulcus. Geodesic distance was calculated on the *fiducial* surface using algorithms in the pycortex (https://gallantlab.github.io/) package (Gao et al., 2015). Then, we averaged across the vertices within each sulcus and tested for a linear relationship between average distance from the central sulcus and myelin content. To take advantage of each subject’s individual data, we built a mixed linear model (random intercepts) in the *lme4* R package, using sulci and hemisphere as explanatory variables to correlate with average myelin content (**Figure 3**).

**Figure 3.**
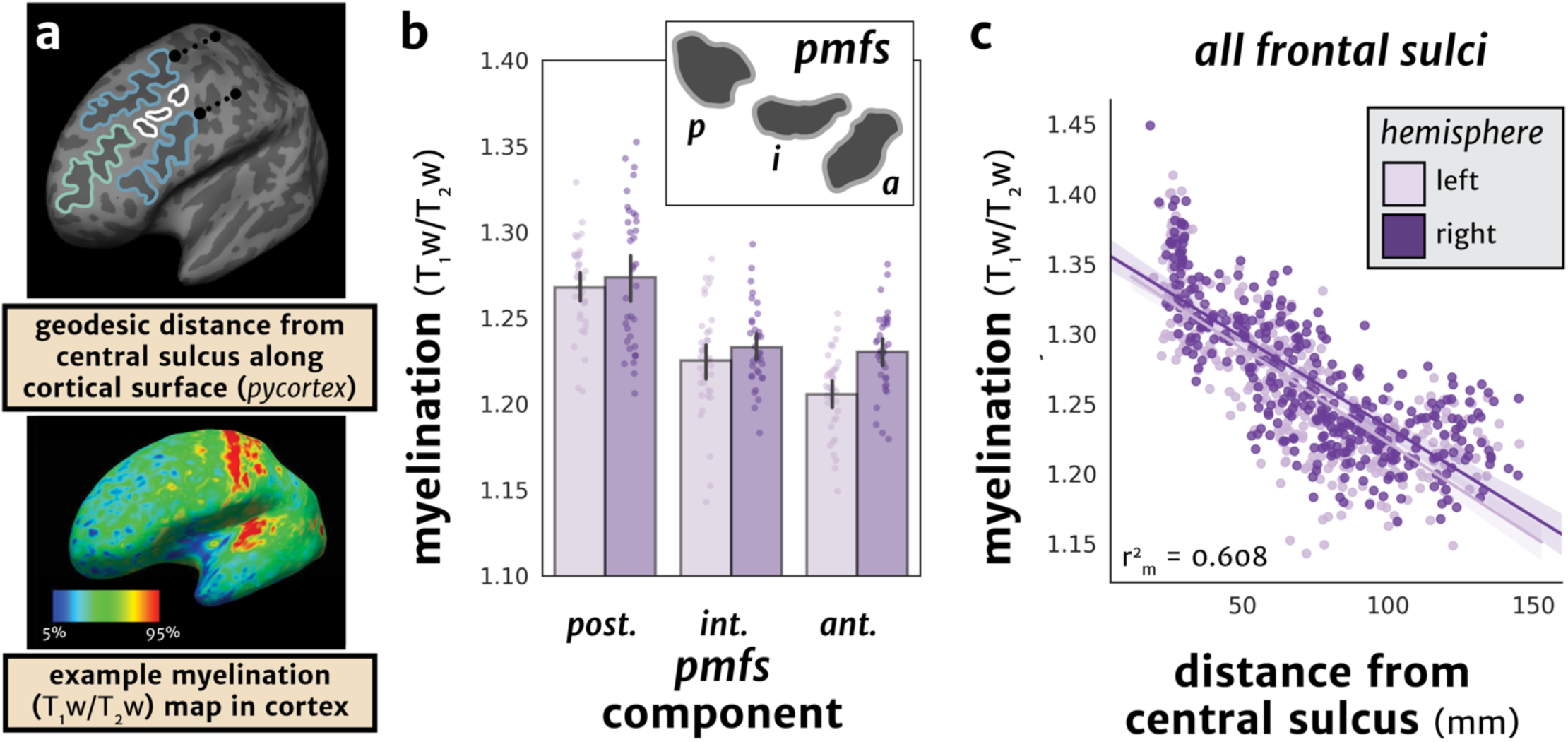
The *pmfs* sulci are anatomically differentiable based on myelin content. (**a**) Top: Schematic of the calculation of geodesic distance along the cortical surface. For each sulcus, the average distance of each vertex from the central sulcus was calculated (dotted black line; **Materials and Methods**). Bottom: an example T_1_w/T_2_w map in an individual subject in which 5-95% percentile of values are depicted. Tissue contrast enhancement (a proxy for myelin) is highest (warm colors) in primary sensorimotor areas. (**b**) Myelination (T_1_w/T_2_w) values plotted for each component of the *pmfs* for each individual subject (N = 36). Bars represent mean ± 95% CI, while each subject is depicted as a circle. Darker shades indicate right hemisphere values, while lighter shades indicate left hemisphere values. The components of the pmfs are differentiable based on myelin content, with a decrease from posterior to anterior across both hemispheres. (**c**) Scatterplot showing the negative relationship between distance from the central sulcus to the mean myelination value for all labeled sulci from each individual (N = 36 subjects). The mixed linear model (**Materials and Methods**) with predictors of distance and hemisphere shows a marginal r^2^ of 60.8%. Scatterplot is bootstrapped at 68% CI for visualization. *Dark purple*: right hemisphere; *Light purple*: left hemisphere.

### Sampling T_1_w/T_2_w myelin index across cortical depths

In order to investigate the microstructural profile of the *pmfs* across cortical layers, we generated nine surfaces from the outermost (*pial*) to the innermost (*white matter)* layers in all of the manually labeled hemispheres using an equivolumetric approach (Waehnert et al., 2014). We implemented the equivolume surface algorithm spanning nine cortical depths with the *surfacetools* Python package that builds on top of FreeSurfer outputs: https://github.com/kwagstyl/surface_tools (Dale et al., 1999). The high-resolution T_1_w/T_2_w volumetric data in each HCP subject’s native anatomical space were then sampled onto each equivolume surface using the FreeSurfer *mri_vol2surf* function to obtain a value of T_1_w/T_2_w at each cortical depth. We compared the mean T_1_w/T_2_w value across depths for each subject in the manually defined *pmfs* components and the surrounding middle frontal gyrus (as defined by FreeSurfer parcellations (Destrieux et al., 2010), but with the *pmfs* components removed). We then conducted a repeated-measures ANOVA followed by post-hoc t-tests at each depth to test for differences between the *pmfs* components and *MFG* in myelin content (**Figure 4**). Tests across each of the nine cortical depths were multiple comparisons corrected at a familywise error (FWE) threshold of *p* = 0.05/9.

**Figure 4.**
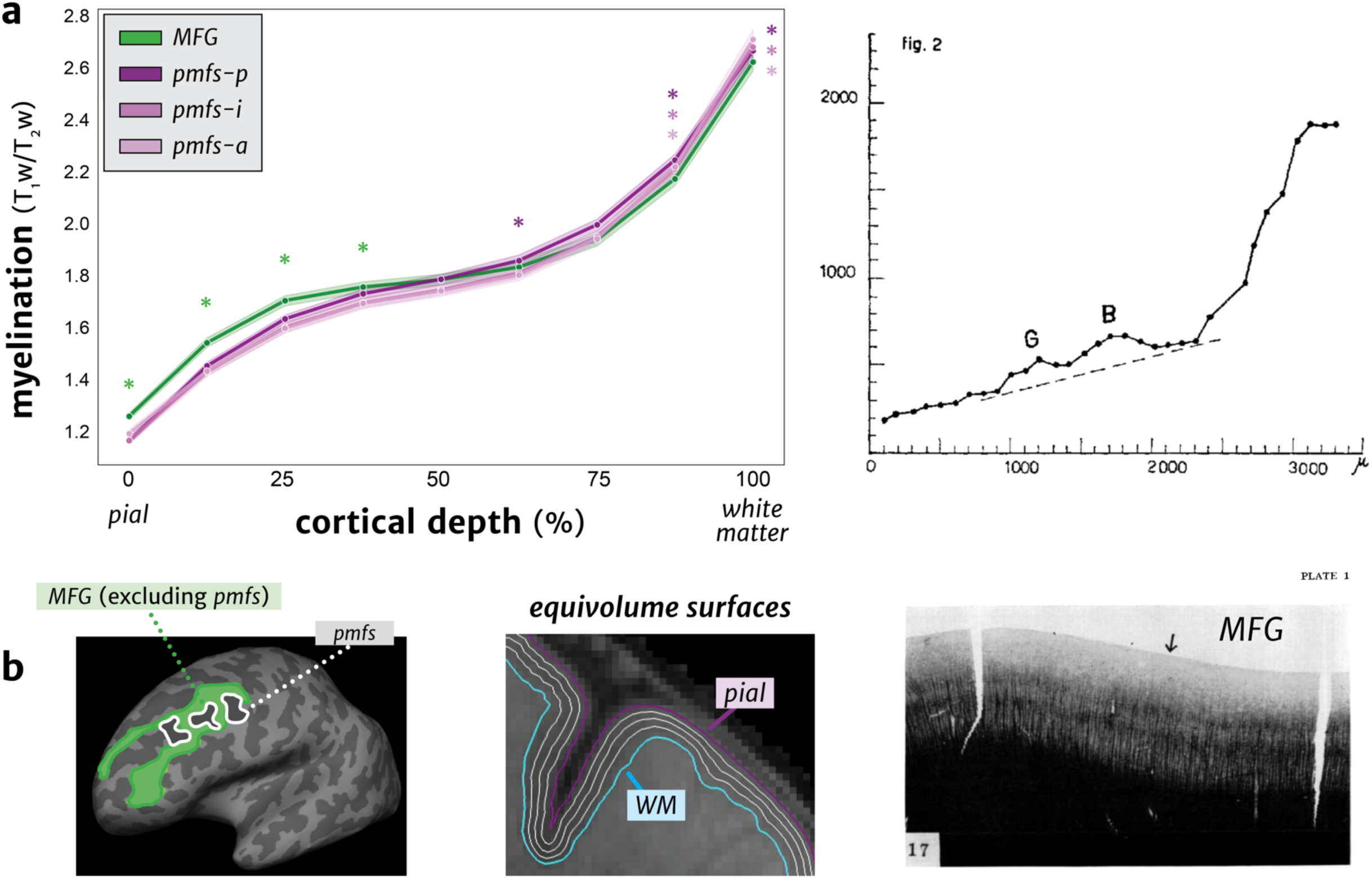
The *pmfs* sulci and middle frontal gyrus have differentiable myelin profiles across cortical depths. (**a**) Left: Tissue contrast enhancement (T_1_w/T_2_w metric, a proxy for myelin) at nine cortical depths, sampled from the outer gray matter (*pial*) to the gray/white matter boundary (*white matter*) using equivolume surfaces (**Materials and Methods**). The *MFG* (excluding the *pmfs*) has higher myelin content than all *pmfs* components in the upper cortical layers, while the *pmfs* components have higher myelin content in deeper layers. Shaded area represents bootstrapped 68% CI across subjects. Green asterisks show significant statistical differences between the MFG and all *pmfs* components (MFG higher), purple asterisks show differences where the *pmfs* components have higher myelin content than the MFG (all tests FWE-corrected at *p* < 0.05/9). Right: Myelinated fiber density (y-axis) profile across cortical depths (x-axis) in post-mortem histological sections of the MFG, adapted from Braitenberg (1962). B: stria of Baillarger. G: stria of Gennari. Similar to our measurements, myelination increases from outer to inner layers within the MFG. (**b**) Left: Individual left hemisphere with the manually defined *pmfs* components (white) and surrounding gyrus, the *MFG*, (green) as defined by FreeSurfer (Destrieux et al., 2010). We excluded the *pmfs* components from the MFG to test for anatomically distinct profiles. Middle: Example equi-volume surfaces at five different cortical depths, from the *pial* to *white matter* surfaces, which were used to sample the T_1_w/T_2_w metric across depths. Right: Myelination stain of a post-mortem histological section of the MFG from Braitenberg (1962). Arrow: Location from which the myelinated fiber density profile in (**a**) was calculated.

**Figure 5.**
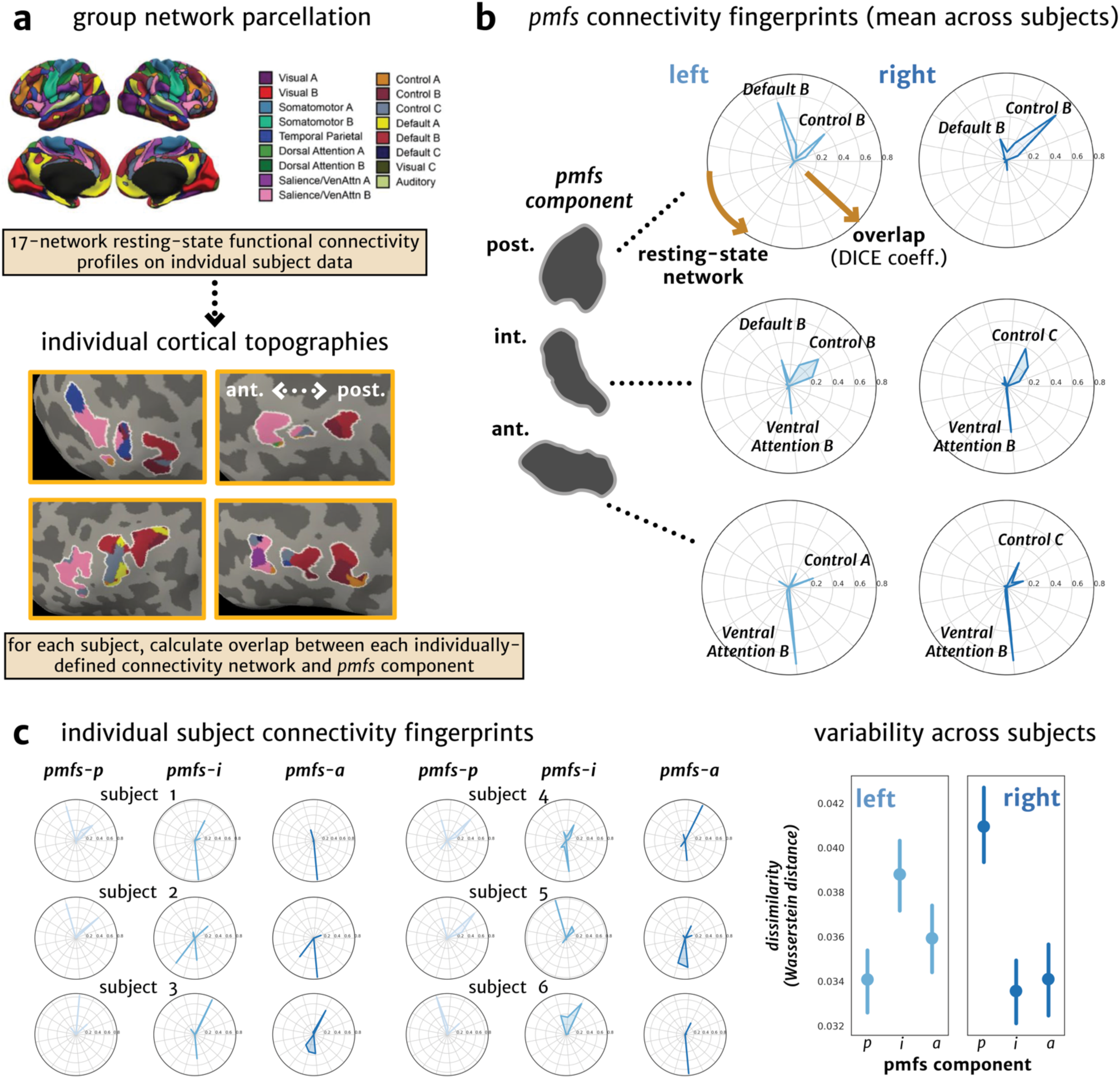
The *pmfs* components are functionally differentiable based on connectivity fingerprints within individual subjects. (**a**) Schematic of how individual-level resting state connectivity profiles were generated in each subject. Resting-state network parcellations for each subject were obtained from a recent study (Kong et al., 2018) in an observer-independent fashion of sulcal definitions in LPFC. Example individual connectivity profiles are shown in four individual subjects, colored according to the group parcellation. The individual connectivity profiles and *pmfs* sulcal definitions were used to calculate the connectivity fingerprint, which represents the overlap of each network within the *pmfs* component of each subject. (**b**) Polar plots showing the mean connectivity fingerprint of the three *pmfs* components (plotted outwards) with each of 17 resting-state functional connectivity networks, across subjects. Resting-state networks with the highest overlap across subjects are labeled. (**c**) Left: Polar plots showing variability among 6 individual subjects. Right: Dissimilarity of the resting-state network fingerprints (variability in the connectivity fingerprint across subjects represented by the Wasserstein distance between unique subject pairs; **Materials and Methods**) are plotted as a function of each *pmfs* component for left and right hemispheres. Error bars represent 68% CI (SEM) across unique subject pairs.

### Cross-validation of sulcal location

In order to quantify the ability to predict the location of each sulcus across subjects, we registered all sulcal labels to a common template surface (*fsaverage*) using cortex-based alignment (Fischl et al., 1999b). Similarity between each transformed individual label and the labels defined on *fsaverage* was calculated via the DICE coefficient, where X and Y are each label:

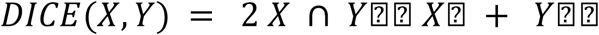

The cortex-based alignment algorithm aligns the surfaces based on sulcal depth and curvature metrics. We use the central sulcus as a proxy noise ceiling measurement for DICE coefficient values from other frontal sulci because it is a large and deep sulcus and is used in the surface registration algorithm that aligns cortical surfaces across subjects (Fischl et al., 1999b).

Sulcal probability maps were calculated to describe the vertices with the highest alignment across subjects for a given sulcus. A map was generated for each sulcus by calculating, at each vertex in the *fsaverage* hemisphere, the number of subjects with that vertex labeled as the given sulcus, divided by the total number of subjects. In order to avoid overlap among sulci, we then constrained the probability maps into *maximum probability maps* (MPMs) by only including vertices where (1) greater than 33% of subjects included the given sulcal label and (2) the sulcus with the highest value of subject overlap was assigned to a given vertex. In a leave-one-subject out cross-validation procedure, we generated probability maps from n = 35 subjects and registered the probability map to the held-out subject’s native cortical surface. This provided a measure of sulcal variability and prediction accuracy (**Figure 7**). This procedure also allows the identification of the *pmfs* sulcal components within held-out individual subjects, reducing the extent of manual labeling necessary to identify this structure. Finally, the MPMs were used when analyzing meta-analytical functional data (described in the section *Cognitive Component Modeling*). The MPMs and code for alignment to new subjects will be available on OSF with the publication of this paper.

**Figure 6.**
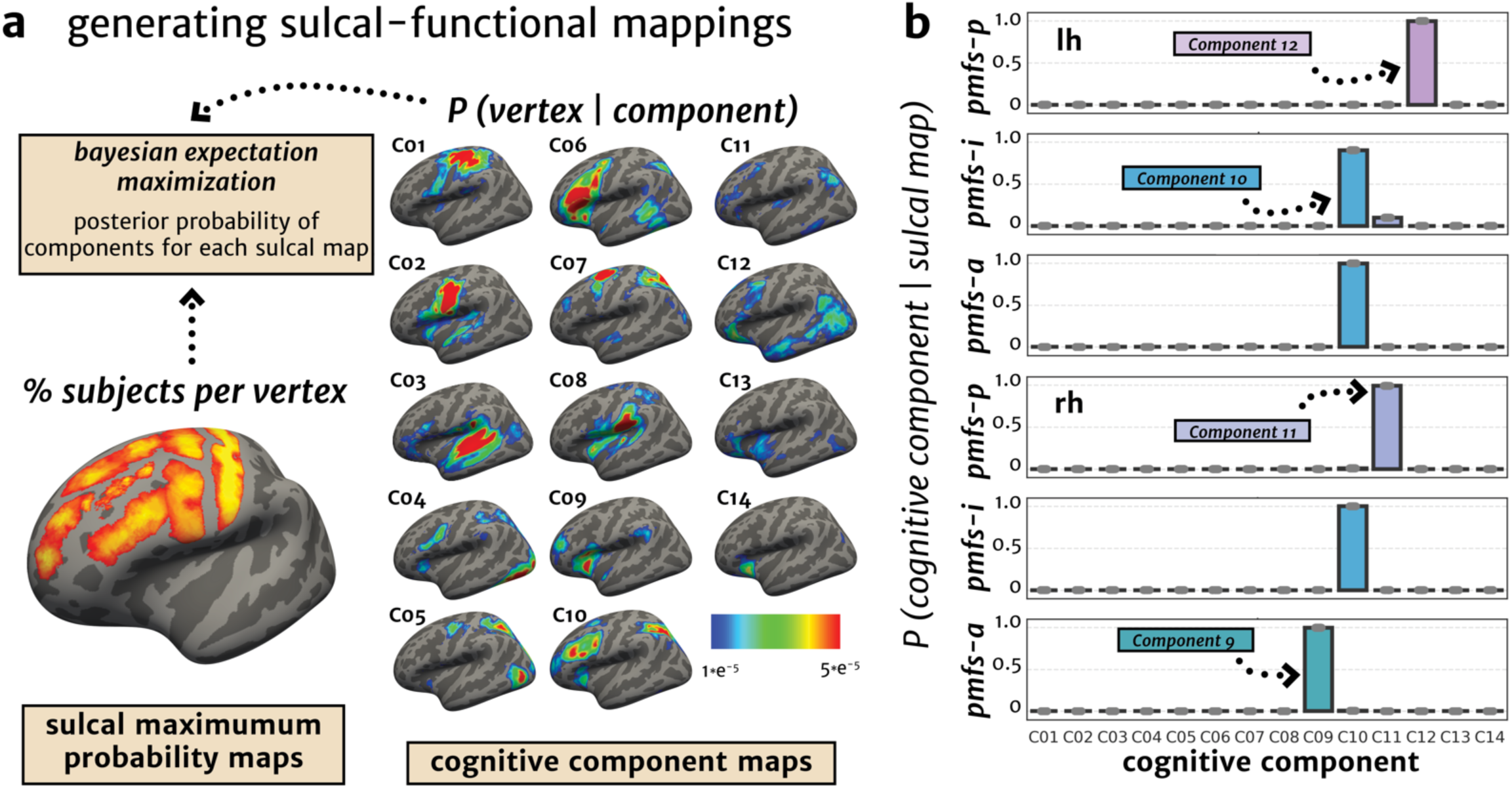
The *pmfs-p, pmfs-i*, and *pmfs-a* are functionally differentiable based on cognitive components: A meta-analysis of fMRI experimental tasks. (**a**) Schematic of analyses linking sulcal probability maps (bottom, left) and cognitive component maps (right) from a meta-analysis of fMRI experimental tasks (Yeo et al., 2015) using an expectation maximization algorithm (**Materials and Methods**). For each *pmfs* component, the algorithm provides a posterior probability for each of 14 cognitive components being associated with the provided sulcal probability map. (**b**) For each *pmfs* component in each hemisphere, the posterior probability for each cognitive component is plotted. This approach reveals that the *pmfs-p* (Component 12, lh; Component 11, rh), *pmfs-i* (Component 10, lh and rh), and *pmfs-a* (Component 10, lh; Component 9, rh; **Materials and Methods**) are functionally dissociable based on meta-analytic data of cognitive task activations. Gray dots indicate individual subject data points when the analysis is performed with individual labels transformed to a template brain, rather than with probability maps.

**Figure 7.**
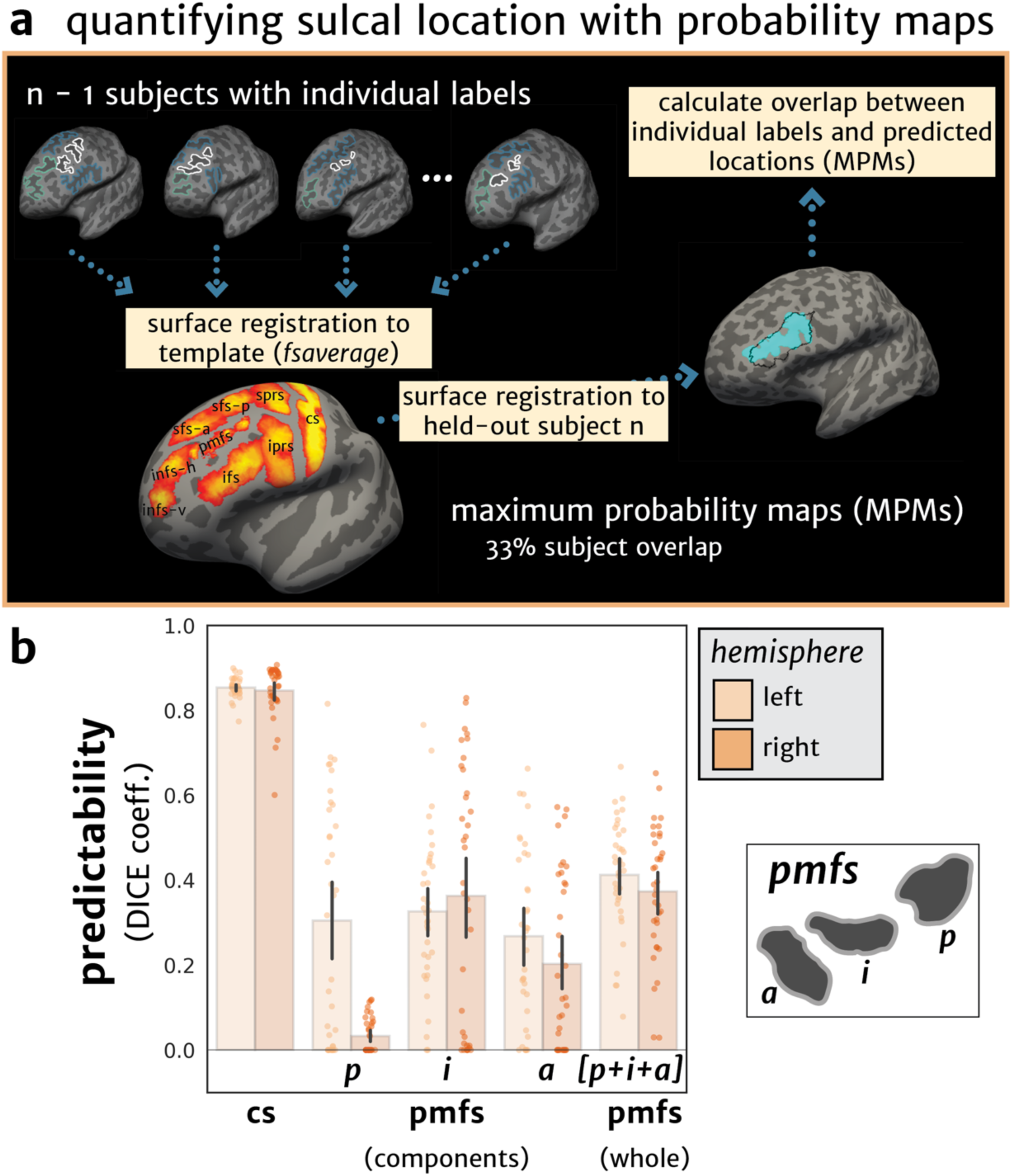
Quantification and prediction of *pmfs-p, pmfs-i, and pmfs-a* within individual hemispheres. (**a**) Procedure to generate sulcal probability maps based on the manual anatomical labeling within each individual subject. Labels from each individual are transformed to a template brain and then projected onto the surface of a held-out individual subject. The overlap between the manual anatomical label and predicted location was then calculated for each iteration across subjects. (**b**) Overlap (DICE coefficient) between predicted and manual location of the *pmfs* components within individual subjects. Prediction is highest when all three components are combined (far right). The central sulcus (*cs*) is included as a noise ceiling for reference, as this landmark is used in the surface registration algorithm that aligns cortical surfaces across subjects.

#### Functional metrics

### Resting-state network topology

In order to calculate if the three *pmfs* sulcal components were functionally distinct from one another, we calculated functional connectivity network profiles for each sulcus. Resting-state network parcellations for each individual subject were used from Kong et al. (2018), who generated individual network definitions by applying a hierarchical Bayesian network algorithm to produce maps for each of 17-networks (Yeo et al., 2011) in individual HCP subjects. These data were calculated in the template HCP *fs_LR 32k* space. We resampled the network profiles for each subject onto the *fsaverage* cortical surface and, then, to each native surface using CBIG tools (https://github.com/ThomasYeoLab/CBIG). We then calculated the overlap of each *pmfs* sulcus in each subject with each of the 17 resting-state networks. We also separated the components of the *pmfs* and tested whether they showed similar or different network connectivity profiles using a 3-way repeated-measures ANOVA (sulcal component x network x hemisphere). Variability across individuals in the network profiles for each *pmfs* component was calculated by generating the Wasserstein metric (Earth Mover’s Distance) between the resting-state network overlap values for each unique subject pair (**Figure 4b**).

### Cognitive component modeling

To further examine if the functional dissociation of the *pmfs-p, pmfs-i, and pmfs-a* is also identifiable using meta-analytic fMRI data, we quantified the overlap between the *maximum probability maps* (MPMs) of each sulcal component and meta-analytic fMRI data from hundreds of experiments aligned to the *fsaverage* surface. Specifically, we quantitatively related the sulcal MPMs to vertex-wise maps for 14 cognitive components, which quantify how each vertex is recruited in a given set of cognitive operations across tasks and experiments (Yeo et al., 2015). We used a Bayesian method of expectation maximization to determine the combination of cognitive components that best fit each sulcal MPM. This resulted in a set of probabilities for each cognitive component for each sulcal map. We tested whether all sulci and the three components of the *pmfs* were distinguishable based upon these cognitive component loadings from a repeated-measures ANOVA (**Figure 5**).

### Statistical methods

All repeated measures ANOVAs (including sphericity correction) and post-hoc t-tests were performed with the *afex* and *emmeans* R packages, imported into Python via *rpy2*. For each repeated measures ANOVA, cortical hemisphere and sulcus were used as within-subject factors. Effect sizes for each main effects and interactions were calculated and reported with the *generalized eta-squared* metric (Fritz et al., 2012). Mixed linear models were implemented in the *lme4* R package. Cortical surface files were loaded in and operated on in Python using the nilearn software: https://nilearn.github.io

## Results

Before conducting our multimodal examination relating morphological features of tertiary sulci to microstructural and functional properties of LPFC, we first had to confront the contradictory nature of historic and modern definitions of sulci within the middle frontal gyrus (MFG). For example, sulcal definitions within the MFG vary in a) their nomenclature, b) the number of sulcal components depicted or acknowledged in schematics, c) the omission or inclusion of sulci within the posterior MFG, and d) the actual empirical data that is included to support the illustration of the sulcal patterning (**Figure 1**). To ameliorate these concerns and to either empirically support or to refute the generality of sulcal definitions within the posterior MFG, we apply a classic, multimodal approach that has been used to distinguish cortical areas from one another in order to determine sulcal definitions in the posterior MFG. Specifically, after identifying each sulcus within the posterior MFG based on a recent proposal in post-mortem human brains (Petrides and Pandya, 2012; Petrides, 2019), we use both anatomical and fMRI data to either support or refute the identification of individual sulci within this cortical expanse. Implementing this two-pronged approach, we first examined if the three components of the posterior middle frontal sulcus (*pmfs*) are consistently identifiable within individual hemispheres. And if so, we then tested if the three *pmfs* components are anatomically and functionally homogenous, or serve to identify anatomical and functional heterogeneity in LPFC. This approach supports the latter in which there are three anatomically and functionally distinct sulci within the posterior MFG: the posterior (*pmfs-p*), intermediate (*pmfs-i*), and anterior (*pmfs-a*) posterior middle frontal sulci.

### Three posterior middle frontal sulci (pmfs) are identifiable within individual subjects and are characteristically shallow

Before examining the sulcal patterning within the posterior MFG, we first identified reliable sulci (**Methods**: *manual sulcal labeling*) surrounding the MFG in both *in vivo* cortical surface reconstructions of MRI data and post-mortem brains (**Figure 2a**). Posteriorly, we identified the central sulcus (*cs*), as well as the superior (*sprs*) and inferior (*iprs*) pre-central sulci. Superiorly, we identified the anterior (*sfs-a*) and posterior (*sfs-p*) superior frontal sulci. Inferiorly, we identified the inferior frontal sulcus (*ifs*). Anteriorly, we identified the horizontal (*imfs-h*) and vertical (*imfs-v*) intermediate frontal sulci. The latter two sulci are consistent with Eberstaller’s classic definition of the middle frontal sulcus, but have since been renamed (**Figure 1**). Within the posterior MFG, we identified three sulci in every hemisphere (N=72). From posterior to anterior, the first sulcus (*pmfs-p*) is positioned immediately anterior to the *sprs* (**Figure 2a, Extended Data Figure 2-1**), and most commonly does not intersect other sulci (see **Extended Data Table 2-1** for a summary of the morphological patterns, or types). The second sulcus (*pmfs-i*) is located immediately anterior to the *pmfs-p*, and typically aligns with the separation between the *sfs-a* and *sfs-p* components. The *pmfs-i* is most often independent (especially in the right hemisphere) or intersects (especially in the left hemisphere) the *pmfs-a*. Finally, the third sulcus (*pmfs-a*) is immediately anterior to the *pmfs-i*, inferior to the *sfs-a*, and posterior to the *imfs-h*. The *pmfs-a* most commonly intersects other sulci in the right hemisphere. Each sulcus is also identifiable within individual *in-vivo* volumetric slices (**Extended Data Figure 2-2**) and in postmortem brains (**Figure 2, Extended Data Figure 2-3**), which indicates that the computational process used to generate the cortical surface reconstruction in the MRI data does not artificially create these sulci within the MFG.

The two most identifying morphological features of the three *pmfs* sulci are their surface area and depth (**Figure 2b**). Each *pmfs* sulcus is of roughly equal surface area (**Figure 2b, Extended Data Table 2-2**), which is smaller than the surface area of the other examined sulci in LPFC (**Figure 2b, Extended Data Table 2-2**). A two-way repeated-measures ANOVA with factors sulcus and hemisphere yielded a main effect of sulcus (*F*(5.78, 202.15) = 384.1, p < 0.001, *η*⍰*G*⍰2⍰ = 0.84) and no main effect of hemisphere (*F*(1, 35) = 0.1, *p* = 0.77). The depth of the three *pmfs* sulci are also the shallowest of the lateral PFC sulci examined (**Figure 2b, Extended Data Table 2-1**). A two-way repeated-measures ANOVA with sulcus and hemisphere as factors yielded a main effect of sulcus (*F*(3.15, 103.84) = 77.7, *p* < 0.001, *η*⍰*G*⍰2⍰ = 0.55), and a main effect of hemisphere (*F(*1, 33) = 20.4, *p* < 0.001, *η*⍰*G*⍰2⍰ = 0.02) in which sulci were deeper in the right compared to the left hemisphere (**Figure 2b, Extended Table 2-2**). Post-hoc tests show that, across hemispheres, the *pmfs-p* is shallower than all other sulci (*p*-values < 0.001, Tukey’s adjustment), and the *pmfs-i* and *pmfs-a* are shallower than all other sulci except for the *imfs-v*. Taken together, three *pmfs* sulci are identifiable in individual hemispheres (**Figure 2, Extended Data Figure 2-1**) and distinguish themselves from other LPFC sulci based on their surface area and shallowness.

### The pmfs-p, pmfs-i, and pmfs-a are anatomically dissociable and reflect a larger rostro-caudal myelination gradient in LPFC

While the *pmfs-p, pmfs-i*, and *pmfs-a* are morphologically distinct from surrounding sulci (**Figure 2**), it is presently unknown if they are anatomically and functionally similar or distinct from one another. To test this, we first extracted and compared average MRI T_1_w/T_2_w ratio values from each sulcus. The T_1_w/T_2_w ratio is a tissue contrast enhancement index that is correlated with myelin content (**Figure 3a**; (Glasser and Van Essen, 2011; Shams et al., 2019). We chose this index because myeloarchitecture is a classic criterion used to separate cortical areas from one another (Vogt and Vogt, 1919; Flechsig, 1920; Hopf, 1956; Dick et al., 2012). A two-way repeated-measures ANOVA with sulcus and hemisphere as factors yielded a main effect of sulcus (*F*(1.76, 61.7) = 85.0, *p* < 0.001, *η*17*G*17217 = 0.39) and a main effect of hemisphere (*F*(1, 35) = 10.5, *p* = 0.003, *η*⍰*G*⍰2⍰ = 0.05) on myelin content, but no sulcus x hemisphere interaction (*F*(1.73, 60.5) = 2.5, *p* = 0.10). The differences in myelin across sulci were driven by the finding that T_1_w/T_2_w decreased from posterior to anterior across hemispheres: *pmfs-p* vs. *pmfs-i*, t(70) = 9.75, *p* < 0.001 (Tukey’s post-hoc), *pmfs-i* vs. *pmfs-a*, t(70) = 2.62, *p* = 0.029, and *pmfs-p* vs. *pmfs-a*, t(70) = 12.37, *p* < 0.001. The right hemisphere also had higher myelin content overall in the *pmfs*, t(35) = 3.25, *p* = 0.003. Accordingly, the three sulci are differentiable based on myelin content in both hemispheres (**Figure 3b**).

The rostro-caudal gradient among the *pmfs-p, pmfs-i*, and *pmfs-a* sulci is embedded within a larger rostro-caudal myelination gradient in lateral PFC. Specifically, modeling T_1_w/T_2_w content across frontal sulci as a function of distance from the central sulcus (**Figure 3c**) using a mixed linear model revealed a significant, negative effect of distance from the central sulcus along the rostral-caudal axis (*β*= −0.001, z = −33.8, *p* < 0.001), with no differences between hemispheres (*β*= −0.003, z = −0.8, *p* = 0.4). Together, our quantifications show that the *pmfs-p, pmfs-i*, and *pmfs-a* are embedded within a larger anatomical and functional hierarchical gradient in LPFC (see **Discussion** for further details).

### The pmfs components show a microstructural profile across cortical layers that is distinct from adjacent gyral areas

Classic and modern findings show that there is generally more intracortical myelin in deeper cortical layers and that the depths of sulci often have less myelinated fibers than gyral crowns (Braitenberg, 1962; Sanides, 1972; Welker, 1990; Annese et al., 2004; Rowley et al., 2015). Building on this work, we sought to calculate microstructural profiles for myelin content across cortical depths for each *pmfs* component, as well as the gyral components of the MFG that surround them (**Figure 4**; **Materials and Methods**). To do this, we implemented equivolume algorithms, which have been used to model cortical laminar organization *in vivo* that corresponds with cyto- and myeloarchitecture in post-mortem histological sections (Waehnert et al., 2014; Paquola et al., 2019).

The MFG and *pmfs* components show distinct microstructural profiles of myelin content across cortical depths. A three-way repeated-measures ANOVA with factors of structure (*pmfs-p, pmfs-i, pmfs-a, MFG*), cortical depth (*0%, 12*.*5%, 25%, 37*.*5%, 50%, 62*.*5%, 75%, 87*.*5%, 100%*), and hemisphere (*left, right*), yields main effects of structure (*F*(2.26, 78.94) = 15.6, *p* < 0.001, *η*⍰*G*⍰2⍰ = 0.007), depth (*F*(1.39, 48.49) = 1849.6, *p* < 0.001, *η*⍰*G*⍰2⍰ = 0.84), and a structure x depth interaction (*F*(6.78, 237.43) = 78.5, *p* < 0.001, *η*⍰*G*⍰2⍰ = 0.02). This interaction between structure and depth did not differ by hemisphere (*F*(4.69, 164.26) = 1.13, *p* = 0.35, *η*⍰*G*⍰2⍰ = 0.02), so subsequent analyses are collapsed across hemispheres. To determine what differences drive the distinct profiles in myelin content across cortical layers between the *pmfs* and MFG, we conducted post-hoc tests at each cortical depth (**Figure 4a**). The MFG has higher myelin content in each of the upper cortical depths (0%, 12.5%, 25%, 37.5%) compared to all of the *pmfs* components (all *p-values* < 0.001, FWE-corrected at *α* = 0.05/9 for the 9 cortical depths). In the middle-to-deep layers (50%, 62.5%), the *pmfs-p* has higher myelin content than either the *pmfs-i* (50%: t(105) = 6.4, *p* < 0.001; 62.5%: t(105) = 7.0, *p* < 0.001) or *pmfs-a* (50%: t(105) = 7.1, *p* < 0.001; 62.5%: t(105) = 8.1, *p* < 0.001), and was even higher than the MFG (50%: t(105) = 0.27, *p* = 0.99; 62.5%: t(105) = 3.7, *p* = 0.002). At the deepest cortical layers, closest to the gray/white matter boundary, all three *pmfs* components show increases in myelin relative to the MFG. Specifically, the *pmfs-a* shows the highest myelin content in the deepest layers, but all three *pmfs* components display higher myelin than the MFG (all *p-values* < 0.001, FWE-corrected at *α* = 0.05/9 for the 9 cortical depths). Altogether, the *pmfs* differs from the MFG in microstructure across cortical layers, with lower myelin content in upper layers, and higher myelin content in deeper layers. This surface-based sampling of cortical depths provides *in vivo* neuroimaging evidence for a microanatomical distinction of the *pmfs* from the surrounding MFG. Further, the depth profiles of T1/T2 within the MFG are similar to classic myeloarchitectural quantifications of the MFG (**Figure 4**).

### The pmfs-p, pmfs-i, and pmfs-a exhibit different characteristic patterns of whole brain functional connectivity

To determine if the *pmfs-p, pmfs-i*, and *pmfs-*a are functionally distinct, we leveraged detailed individual functional parcellations of the entire cerebral cortex based on functional connectivity from a recently published study (Kong et al., 2018) (**Figure 5a**). Importantly, this parcellation was conducted blind to both cortical folding and our sulcal definitions. Within each hemisphere in the same subjects in which we generated manual sulcal labels, we generated a functional connectivity network profile (which we refer to as a “connectivity fingerprint”). For each sulcal component, we calculated the overlap between 17 functional networks (on the native hemisphere, based on the DICE coefficient; **Materials and Methods**). This technique generated a cortical topography reflective of the whole-brain connectivity patterns for each sulcal component (**Figure 5a, bottom**), and can be interpreted similarly to other studies of functional network variations (Gordon et al., 2017; Seitzman et al., 2019), as a trait-like connectivity profile for each *pmfs* component within each participant.

Our approach demonstrated that the *pmfs-p, pmfs-i*, and *pmfs-a* have different connectivity fingerprints and thus, are functionally dissociable. Average connectivity fingerprints across subjects are illustrated in **Figure 5b**. A repeated-measures ANOVA with sulcal component (*pmfs-p, pmfs-i, pmfs-a*), hemisphere (left, right), and network yielded a significant component x network interaction (*F*(32, 1120) = 45.2, *p* < 0.001, *η*⍰*G*⍰2⍰ = 0.29), as well as a component x network x hemisphere interaction (*F*(32, 1120) = 5.26, *p* < 0.001, *η*⍰*G*⍰2⍰ = 0.040) (**Figure 5b**). In each hemisphere, there is a component x network interaction (left: *F*(32, 1120) = 29.4, *p* < 0.001, *η*⍰*G*⍰2⍰ = 0.35, right: *F*(32, 1120) = 23.2, *p* < 0.001, *η*⍰*G*⍰2⍰ = 0.27) in which the difference between hemispheres is driven by the *pmfs-p* connectivity fingerprint. Specifically, the *pmfs-p* overlaps most with the default mode network in the left hemisphere and the cognitive control network in the right hemisphere.

Additionally, there are also individual and hemispheric differences in the connectivity fingerprint of each *pmfs* component at the single subject level (**Figure 5c**; **Extended Data Figure 5-1**). To characterize individual differences, we built on work showing network connectivity variations across individuals (Kong et al., 2018; Seitzman et al., 2019) by relating this connectivity variability to individual anatomical landmarks in LPFC. We quantified connectivity fingerprint variability by measuring the pairwise Wasserstein distance between the connectivity profiles for all unique subject pairs for each sulcal component, in which higher distance indicates decreased similarity, and therefore greater variability (see **Materials and Methods**). This approach quantifies how variable the pattern of network overlap (connectivity fingerprint) is across individuals for each *pmfs* component (**Figure 5c**, right). In the right hemisphere, the *pmfs-p* showed the most variable network profile across all unique subject pairs (*pmfs-p* vs. *pmfs-i*, Wilcoxson-Signed rank test, *W* = 7.2×10^4^, *p* < 0.001, *pmfs-p* vs. *pmfs-a, W* = 7.4×10^4^, *p* < 0.001), while the *pmfs-i* was most variable in the left hemisphere (*pmfs-i* vs. *pmfs-a, W* = 8.8×10^4^, *p* = 0.014, *pmfs-i* vs. *pmfs-p, W* = 8.0×10^4^, *p* < 0.001). This analysis suggests that the right *pmfs-p* and left *pmfs-i* mark regions of LPFC with particularly high levels of individual differences in functional connectivity profiles, providing an anatomical substrate for network connectivity differences across individuals.

### The pmfs-p, pmfs-i, and pmfs-a are functionally dissociable: Meta-analyses across 83 experimental task categories

We next tested if the dissociation of functional networks between the *pmfs-p, pmfs-i*, and *pmfs-a* identified in individual subjects (**Figure 5**) can also be observed in meta-analytic analyses of functional activation data at the group-level. That is, do the components of the *pmfs* show a functional dissociation of engagement over a wide array of cognitive operations? To test for different patterns of functional activations across tasks, we generated sulcal probability maps on a template cortical surface (**Figure 6a**, bottom left). Analogous to probabilistic maps for functional regions (Wang et al., 2015), the maps provide a vertex-wise measure of anatomical overlap across individuals for all 13 sulci in LPFC. As the *pmfs* components disappear on average templates (**Figure 1**), these probabilistic maps are independent of the sulcal patterning of the template itself, which merely serves as a cortical surface independent of each individual cortical surface. We then compared these sulcal probability maps to 14 probabilistic “cognitive component” maps derived from an author-topic model of meta-analytic activation data across 83 experimental task categories (Yeo et al., 2015).

The cognitive component model links patterns of brain activity to behavioral tasks via latent components representing putative functional subsystems (Yeo et al., 2015). Each cognitive component map (which was calculated on the same template cortical surface) provides the probability that a given voxel will be activated by each of the 14 components (across all 83 tasks). We then used an expectation maximization algorithm (via posterior probability, **Materials and Methods**) to relate brain activity in each sulcal probability map to each cognitive component (**Figure 6a**, right). Importantly, when calculating the posterior probabilities, we implemented a leave-one-subject-out cross-validation procedure when constructing the sulcal probability maps in order to assess variability in the generated posterior probabilities for each cognitive component (**Fig. 6b**). To indicate feasibility of this approach, the somato-motor components of the cognitive component map (C01, C02) align most highly with the central sulcus (**Extended Data Figure 6-1**) as one would expect, which shows the ability of this method to measure structural-functional correspondences at the meta-analytic level.

This approach further reveals that the *pmfs-p, pmfs-i*, and *pmfs-a* are functionally dissociable based on meta-analytic data of cognitive task activations. In the right hemisphere, the *pmfs-p, pmfs-i*, and *pmfs-a* showed distinct probabilities for separate cognitive components: 1) the *pmfs-p* loaded onto a default mode component (C11), 2) the *pmfs-i* loaded onto an executive function component (C10), and 3) the *pmfs-a* loaded onto an inhibitory control component (C09). In the left hemisphere, the *pmfs-a* and *pmfs-i* both loaded onto an executive function (C10) component, while the *pmfs-p* loaded onto an emotional processing/episodic memory component (C12). Like our individual subject analyses, there were also hemispheric differences: the cognitive components overlapping the most with the *pmfs-a* and *pmfs-p* differed between the two hemispheres. The *pmfs-p* loaded onto an emotional processing/episodic memory component in the left hemisphere (**Figure 6b**, top row) and a default mode component in the right hemisphere (**Figure 6b**, fourth row), while the *pmfs-a* loaded onto an executive function component in the left hemisphere (**Figure 6b**, third row) and an inhibitory control component in the right hemisphere (**Figure 6b**, bottom row).

### Extensive individual differences in the location of the pmfs across individuals

Although the three *pmfs* components are prominent within each hemisphere, there is extensive individual variability in the precise location of each sulcal component within the posterior MFG. To determine how well the probability maps could predict the location of the *pmfs-p, pmfs-i*, and *pmfs-a* within *individual* hemispheres, we used a cross-validated approach, iteratively leaving out one subject from the calculation of probability maps (**Figure 7a**). Then, the maximum probability maps (MPMs) were projected to the held-out individual’s native cortical surface to calculate the overlap between the manually identified and probabilistically identified sulcal locations. This procedure resulted in a measure of location variability for each sulcal component (**Figure 7b**). For these calculations, we used the *central sulcus* (CS) as a noise ceiling (left: *cs* = 0.85 ± 0.02; right: *cs* = 0.85 ± 0.06) as it is a) considered very stable across individuals (see **Materials and Methods**) and b) used in the cortex-based alignment procedure (Fischl et al., 1999b).

The *pmfs* components exhibited significant variability in sulcal location across subjects (left: *pmfs-p* = 0.30 ± 0.28, *pmfs-i* = 0.32 ± 0.18, *pmfs-a* = 0.27 ± 0.20; right: *pmfs-p* = 0.03 ± 0.04, *pmfs-i* = 0.37 ± 0.18, *pmfs-a* = 0.20 ± 0.20). A 2-way repeated-measures ANOVA with *pmfs* sulcal component (*pmfs-p, pms-i, pmfs-a*) and hemisphere (*right, left*) revealed a sulcus x hemisphere interaction (*F*(1.84, 64.47) = 9.52, *p* < 0.001, *η*17*G*17217 = 0.08) driven by the finding that the *pmfs-p* is highly variable across subjects, resulting in very little predictability in the right hemisphere (**Figure 7b**). When using all three *pmfs* components together, prediction is more robust (left: *pmfs* = 0.41 ± 0.13; right: *pmfs* = 0.37 ± 0.15), but still much lower than the predictability of the CS and also lower than prediction performance for all other LPFC sulci quantified in the present study (**Extended Data Figure 7-1**). These results demonstrate that although the *pmfs* is prominent within each individual (**Extended Data Figure 2-1**), the location of each *pmfs* component is variable across individuals, which provides empirical support for the historical confusion regarding its identification and labeling (**Figure 1**).

## Discussion

Here, we examined the relationship between cortical anatomy and function in human lateral prefrontal cortex (LPFC) with a two-pronged approach. First, we tested if a recent novel proposal of tertiary sulci in human LPFC in post-mortem brains (Petrides and Pandya, 2012; Petrides, 2019) could be applied to sulcal definitions of LPFC within *in-vivo* cortical surface reconstructions. Second, we tested if these tertiary LPFC sulcal definitions could serve as a meso-scale link between microstructural and functional properties of human LPFC in individual participants. For the first time (to our knowledge), this two-pronged and multi-modal approach revealed that the posterior middle frontal sulcus (*pmfs*) serves as a meso-scale link between myelin content and functional connectivity in individual subjects. Specifically, the *pmfs* is a characteristically shallow tertiary sulcus with three components that differ in their myelin content, resting state connectivity profiles, and engagement across meta-analyses of 83 cognitive tasks. We first discuss how these findings suggest modern empirical support for a classic, yet largely unconsidered, anatomical theory (Sanides, 1962, 1964), as well as a recent cognitive neuroscience theory proposing a functional hierarchy in LPFC (Koechlin and Summerfield, 2007; Badre and D’Esposito, 2009; Badre and Nee, 2018). We end by discussing a growing need for computational tools that automatically define tertiary sulci throughout cortex.

The anatomical-functional coupling in LPFC identified here is quite surprising considering the widespread literature providing little support for fine-grained anatomical-functional coupling in this cortical expanse and in association cortices more broadly when conducting traditional group-analyses (Paquola et al., 2019; Vazquez-Rodriguez et al., 2019). Indeed, cortical folding patterns relative to the location of anatomical, functional, or multimodal transitions are considered “imperfectly correlated” (Welker, 1990; Glasser et al., 2016) in association cortices and especially in LPFC (Van Essen et al., 2012; Caspers et al., 2013b; Robinson et al., 2014; Coalson et al., 2018). Contrary to these previous findings that did not consider tertiary sulci, the present findings appear to support a classic, yet largely unconsidered theory proposed by Sanides (1962, 1964) that tertiary sulci are potentially meaningful anatomical and functional landmarks in association cortices – and in particular, in LPFC. Specifically, Sanides proposed that because tertiary sulci emerge late in gestation and exhibit a protracted postnatal development, they likely serve as functional and architectonic landmarks in human association cortices, which also exhibit a protracted postnatal development. He further proposed that the late emergence and continued postnatal morphological development of tertiary sulci is likely related to protracted cognitive skills thought to be associated with LPFC such as sustained attention and executive functioning (Sanides, 1964). Interestingly, identifying the three *pmfs* components quantified in the present study in his classic images shows myeloarchitectonic gradations among five areas in LPFC (**Figure 8a**). Linking these data to recent modern multi-modal parcellations of the human cerebral cortex (Glasser et al., 2016) shows that the *pmfs* components likely serve as boundaries among a series of cortical areas, which can be addressed in future research in individual participants (**Figure 8b**).

**Figure 8.**
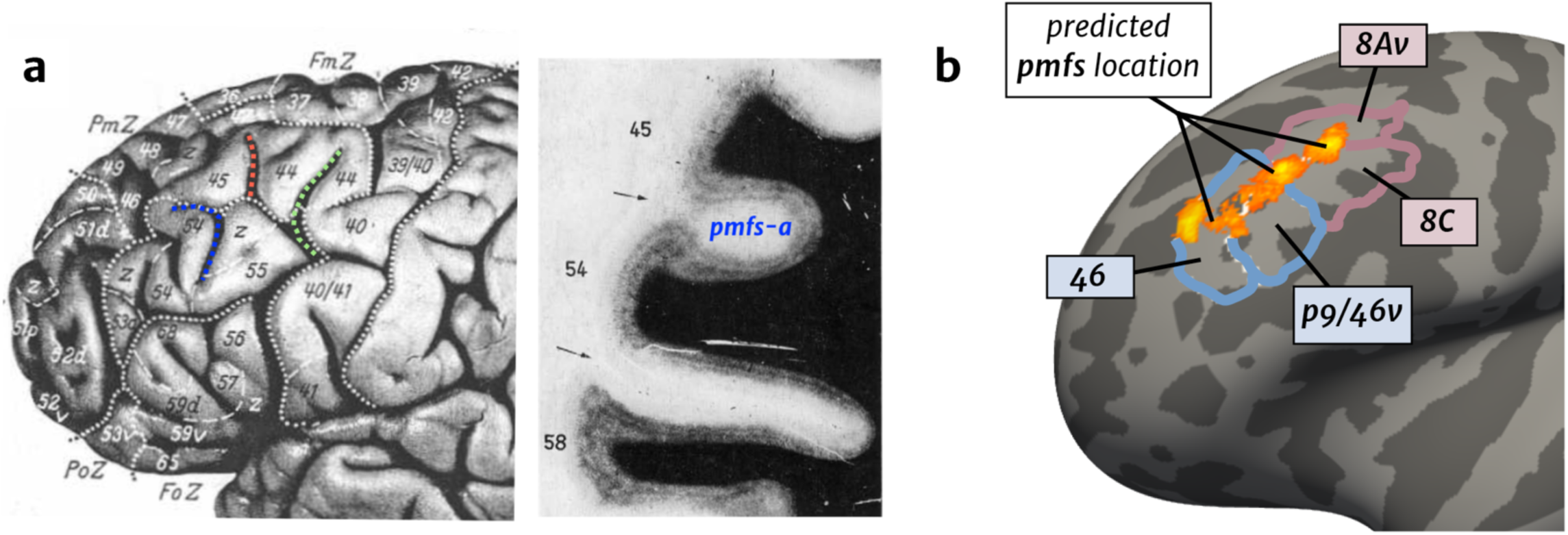
Linking the past to the present: Myelination gradients, multimodal areas, and the *pmfs*. (**a**) Left: Photograph of a left hemisphere from Sanides (1962). Numbers indicate cortical areas differing in myeloarchitecture. Dotted white lines: Sulcal boundaries as defined by Sanides. Dotted colored lines: *pmfs-p* (green), *pmfs-i* (red), and *pmfs-a* (blue) based on modern definitions used in the present study. Identifying *pmfs* components in Sanides’ classic images shows that he identified myeloarchitectonic gradations within *pmfs* components, which is consistent with the present measurements. Gradations occurred in superior-inferior, as well as anterior-posterior dimensions. In the inferior portion of the *pmfs-p* (green), there is an anterior-posterior transition between areas 40 and 55. In the *pmfs-i* (red), there are two transitions: (i) a superior-inferior transition between areas 44 and a transition zone to area 55, and (ii) an anterior-posterior transition between areas 44 and 45. In the *pmfs-a*, there is a transition between areas 45 and 54. Right: Myelination stain of a histological section (coronal orientation) from Sanides (1962). Arrows indicate boundaries between labeled myeloarchitectonic areas (numbers). *pmfs-a* is labeled to help the reader link the myelination stain to the image at left. The reader can appreciate the shallowness of the pmfs-a relative to the sulcus (the IFS) between areas 54 and 58, which is also consistent with our measurements (**Figure 2**). (**b**) Maximum probability maps (thresholded at 33% overlap across subjects) for the *pmfs-p, pmfs-i*, and *pmfs-a* are shown on the Freesurfer average template (left hemisphere). The probability maps are shown relative to four areas from a multi-modal cortical parcellation based on structural and functional MRI data (Glasser et al., 2016). The *pmfs-a* appears to denote the dorsal to ventral transition between areas 46 and p9/46v in anterior LPFC, while *pmfs-p* appears to denote the dorsal to ventral transition between areas 8Av and 8C in posterior LPFC. Future research will be sure to test this hypothesis at the individual subject level.

In addition to supporting Sanides’ classic anatomical theory, the present data demonstrated that the three *pmfs* components exhibit different resting-state intrinsic connectivity profiles along a rostral-caudal axis, which builds on previous work also supporting a functional hierarchy along a rostral-caudal axis of LPFC. Further consistent with this hierarchy, evidence from neuroimaging, lesion, and electrocorticography studies indicate that this proposed rostral-caudal axis of LPFC is also related to levels of temporal and cognitive abstraction. That is, more anterior LPFC cortical regions are more highly engaged in tasks with higher abstract complexity (Koechlin et al., 2003; Koechlin and Summerfield, 2007; Voytek et al., 2015; Mansouri et al., 2017). While there is axonal tracing data in non-human primates suggesting an anatomical basis for such a hierarchical organization (Goulas et al., 2014; Goulas et al., 2019), the present findings provide new evidence for anatomically and functionally dissociable sulcal components in LPFC that also support a hierarchical organization within individuals. Future work leveraging finer-scale multimodal and microanatomical data from individual human brains will be critical for uncovering anatomical and functional properties of LPFC across spatial and temporal scales that may further support the proposed functional rostral-caudal hierarchy of human LPFC.

Together, the culmination of present and previous findings suggest that tertiary sulci are landmarks in human ventral temporal cortex (Nasr et al., 2011; Caspers et al., 2013a; Weiner et al., 2014; Lorenz et al., 2017), medial PFC (Amiez et al., 2019; Lopez-Persem et al., 2019), and now, LPFC. This begs the question: How many other tertiary sulci serve as cortical landmarks? We stress that it is unlikely that all tertiary sulci will serve as cortical landmarks, since neuroanatomists have known for over a century that not all sulci function as cortical landmarks (Smith, 1907; Bailey and Bonin, 1951; Ono et al., 1990; Welker, 1990; Van Essen et al., 2019). Nonetheless, this does not preclude the importance of future studies identifying which tertiary sulci are architectonic, functional, behavioral, or multimodal landmarks – not only in healthy young adults as examined here, but also in developmental (Voorhies et al., 2020) and clinical (Brun et al., 2016; Parker, 2019) cohorts.

Such an exercise of carefully examining the relationship among tertiary sulci and multiple types of anatomical, functional, and behavioral data in individual subjects will require new neuroimaging tools to automatically identify tertiary sulci throughout human cortex. For instance, most neuroimaging software packages are only capable of automatically defining ∼30-35 primary and secondary sulci in a given hemisphere (Destrieux et al., 2010). Current estimates approximate 110 sulci in each hemisphere when considering tertiary sulci (Petrides, 2019). Thus, studies in the immediate future will still require the manual identification of tertiary sulci, which is labor intensive and requires expertise^1^. For example, the present study required manual definitions of 936 sulci in 72 hemispheres. While 72 is a large sample size compared to other labor-intensive anatomical studies in which 20 hemispheres is considered sufficient to encapsulate individual differences (Amunts and Zilles, 2015; Amunts et al., 2020), 2400 hemispheres are available just from the HCP alone. Defining tertiary sulci in only the LPFC of every HCP participant would require ∼26,400 manual definitions, while defining all tertiary sulci in the entire HCP dataset would require over a quarter of a million (∼256,800) manual definitions. Consequently, manual identification of tertiary sulci will continue to limit sample sizes in immediate future studies until new automated methods are generated^2^.

In the interim, we sought to leverage the anatomical labeling in this study to aid the field in the identification of sulcal landmarks in LPFC. The probability maps of sulcal locations in the present study are openly available and may be transformed to held-out individual brains (**Figure 6**). Accordingly, manual identification of these landmarks within individuals is greatly aided, allowing future studies to apply these tools to identify LPFC tertiary in individual subjects, including those from various groups such as patient or developmental cohorts. Because smaller tertiary sulci in association cortex are the latest sulcal indentations to develop (Welker, 1990; Zilles et al., 2013), their anatomical trajectories and properties likely relate to the development of cognitive abilities associated with the LPFC and other association areas as Sanides hypothesized, which recent ongoing work supports (Voorhies et al., 2020). Moving forward, we hope to leverage the manual labeling performed here to develop better automated algorithms for sulcal labeling within individuals. Future work using deep learning algorithms data may help to identify tertiary structures in novel brains without manual labeling or intervention (Borne et al., 2020; Hao et al., 2020). Such automated tools have translational applications as tertiary sulci are largely hominoid-specific structures (Amiez et al., 2019) located in association cortices associated with pathology in many neurological disorders. Thus, morphological features of these under-studied neuroanatomical structures may be useful clinical biomarkers for future diagnostic purposes. To begin to achieve this goal and to aid the field, we share our probabilistic maps of LPFC tertiary sulci with the publication of this paper.

## Data availability

The raw data in the present work is publicly available from the Human Connectome Project (HCP): <https://www.humanconnectome.org/study/hcp-young-adult/document/1200-subjects-datarelease)/ The HCP dataset and processing are described in previous publications (Glasser et al., 2013; Glasser et al., 2016). The probability maps for LPFC sulcal definitions along with anatomical statistics for each manually labeled sulcus and analysis code will be freely available with the publication of the paper.

## Author Contributions

Manual anatomical labeling: J.A.M., W.V., K.S.W. Data analysis and interpretation of the data: J.A.M., D.J.L., K.S.W. Drafting paper: J.A.M, K.S.W. Revising paper: J.A.M, W.V., D.J.L., M.D., K.S.W. Supervision and study conceptualization: J.A.M., M.D, K.S.W.

## Acknowledgments

This work was supported by (1) start-up funds provided by the University of California, Berkeley and the Helen Wills Neuroscience Institute (K.S.W.); (2) NIH grant RO1 MH63901 (J.A.M., M.D.). We also thank Jewelia Yao for help with the manual definition of LPFC sulci.

## Extended Data

**Extended Data Figure 2-1.**
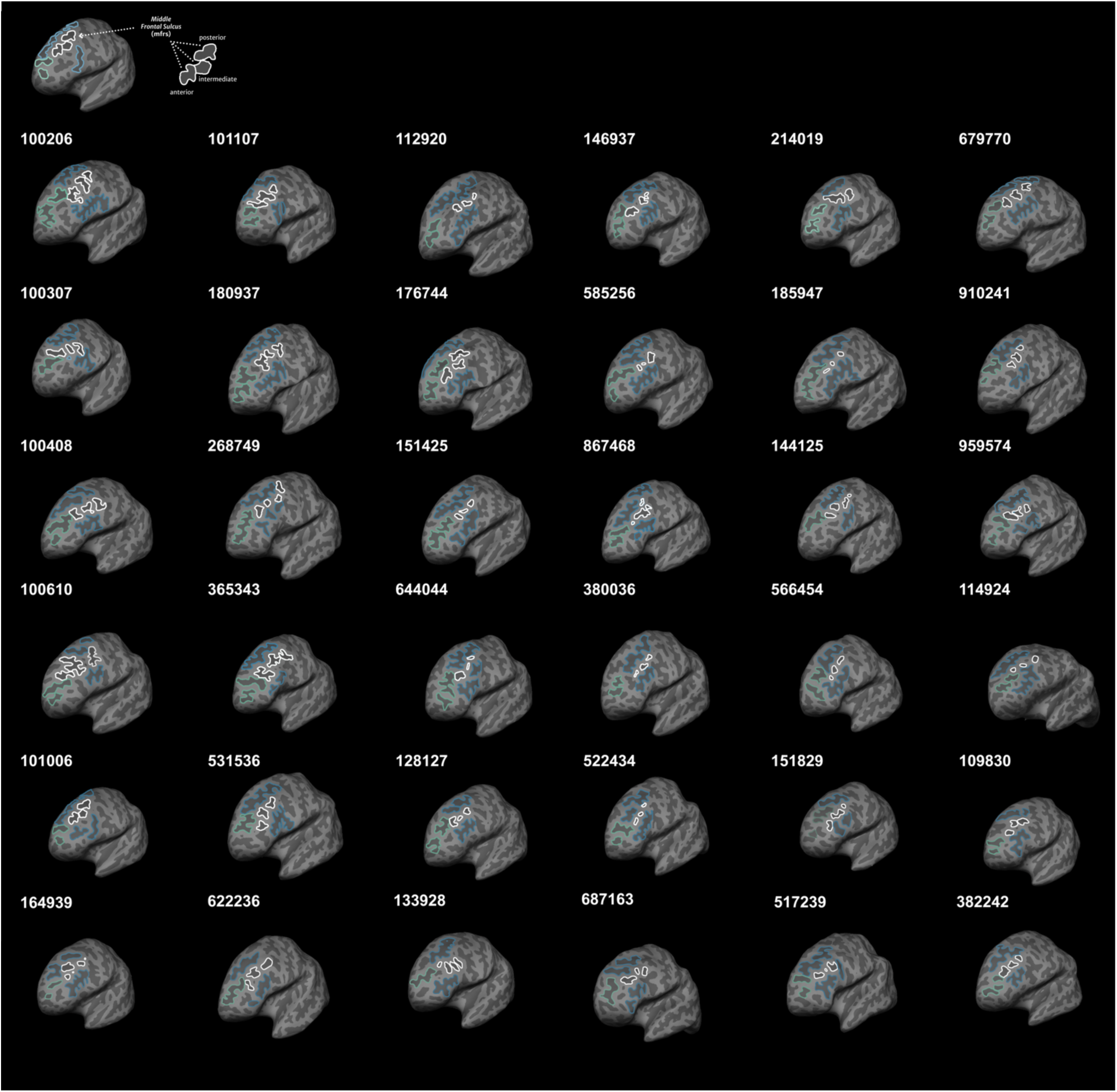
Individual labeling of the *pmfs* in all subjects. As in Figure 1, the posterior middle frontal sulcus (*pmfs*) and its three components are outlined in white on the individual inflated cortical surface of each subject. For reference, the large superior (*sfs*) and inferior (*ifs*) frontal sulci are also outlined, in blue, along with the horizontal (*imfs-h*) and vertical (*imfs-v*) intermediate frontal sulci, in green.

**Extended Data Figure 2-2.**
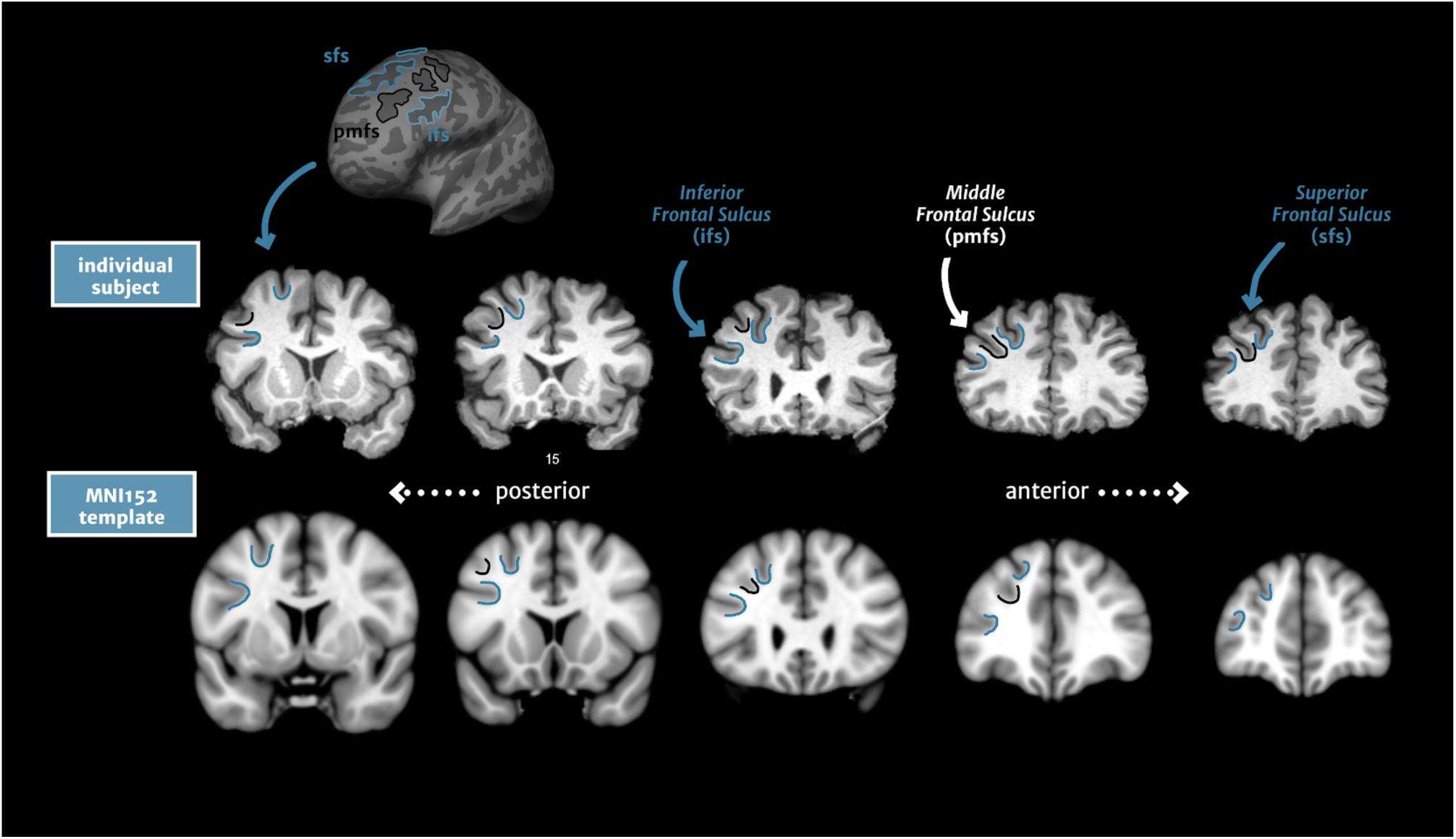
Identification of the *pmfs* on coronal MRI sections. The *pmfs* is easily identifiable on T1 volume MR images. Top: The *pmfs, sfs*, and *ifs* are clearly visible in coronal sections from an individual subject. Bottom: These three landmarks are also identifiable on the MNI152 template brain, although the *pmfs* is much less pronounced. The fact that the *pmfs* becomes obscured on an average volume, is consistent with the fact that it is obscured when averaging cortical surfaces (**Fig. 1**).

**Extended Data Figure 2-3.**
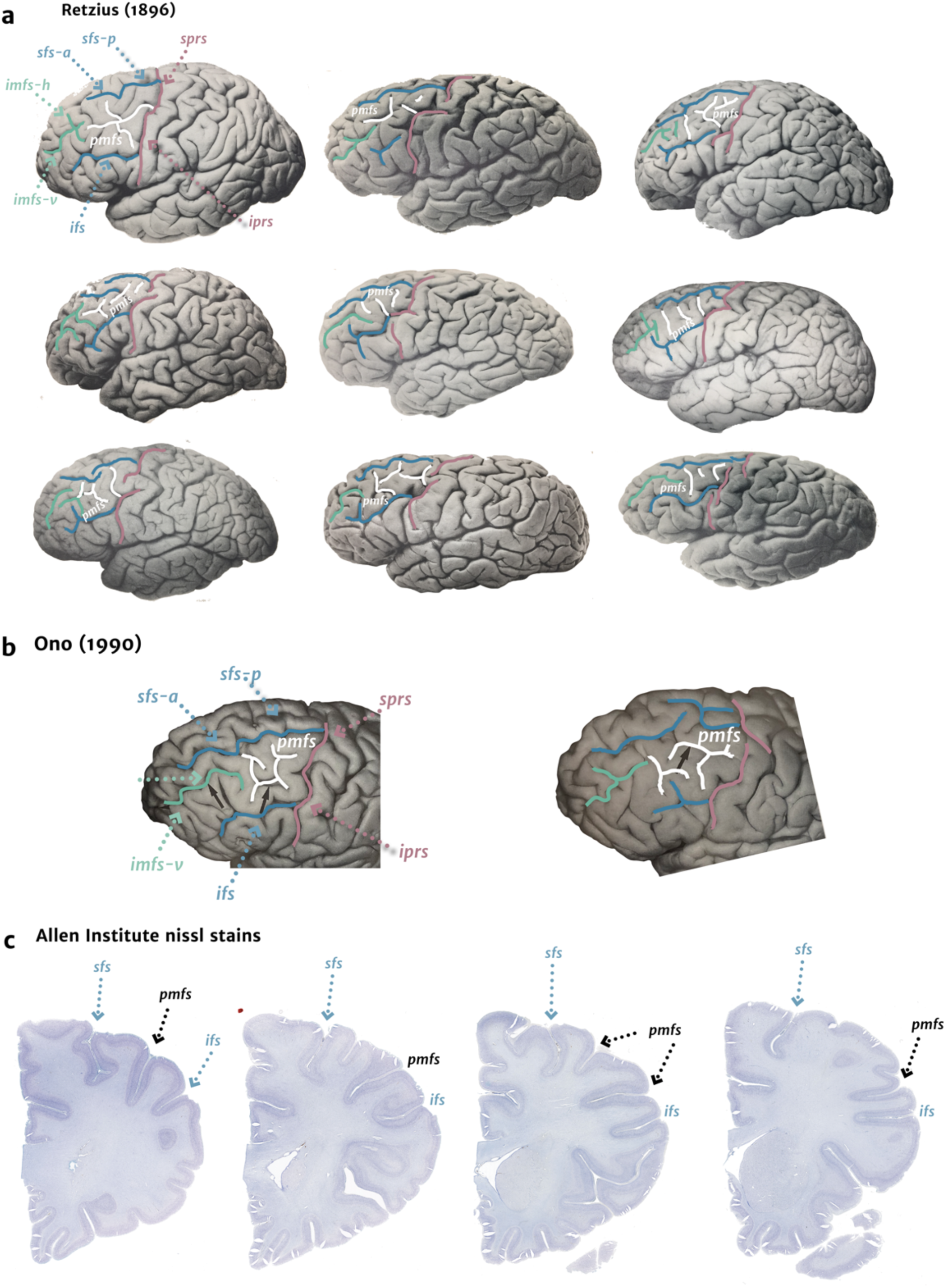
Identification of the *pmfs* in post-mortem hemispheres and histological slices. While our lateral PFC sulcal definition of the pmfs relative to surrounding sulci is guided by data from post-mortem brains (Petrides and Pandya, 2012) and we can easily identify these sulci in in-vivo data, we also applied this sulcal definition to post-mortem brains from classic and modern atlases. (**a**) The lateral PFC sulci delineated on 10 hemispheres from a classic neuroanatomical atlas (Retzius, 1896). The *pmfs* is clearly present between the *sfs* and *ifs* within the MFG. (**b**) The pmfs can also be identified on a more recent atlas of sulcal morphology. In this work (Ono et al., 1990), the sulci defined as the *pmfs* were considered as the posterior end of the *imfs* by the authors. However, our present quantifications show that this sulcus is distinct from the *imfs*, with different morphological and functional features. (**c**) Post-mortem histological slices with Nissl staining from the Allen Institute (*images 14-17* at https://atlas.brain-map.org/atlas?atlas=138322605) showing that the *pmfs* is situated in between the *sfs* and *ifs*.

**Extended Data Table 2-1.**
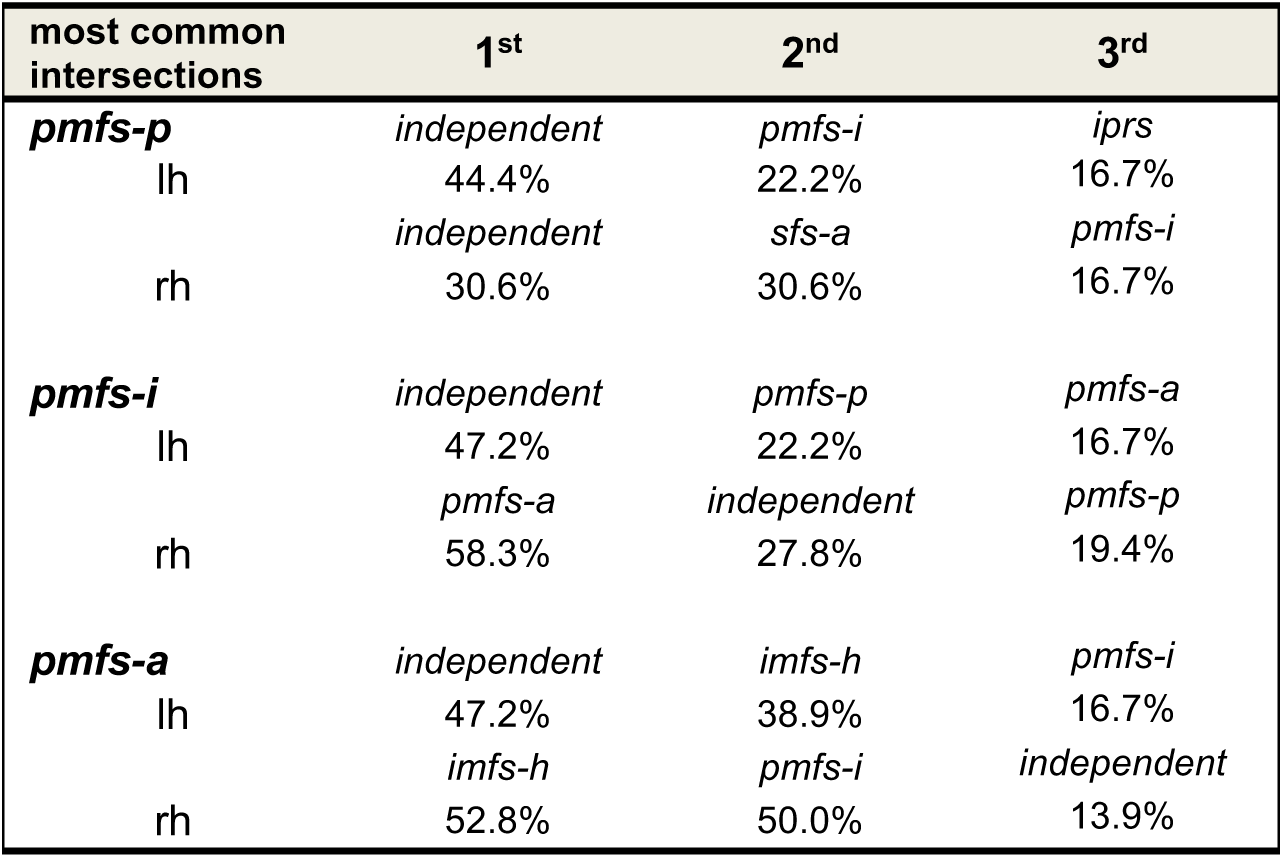
Most common intersections of the *pmfs* components (morphological types).

**Extended Data Table 2-2.**
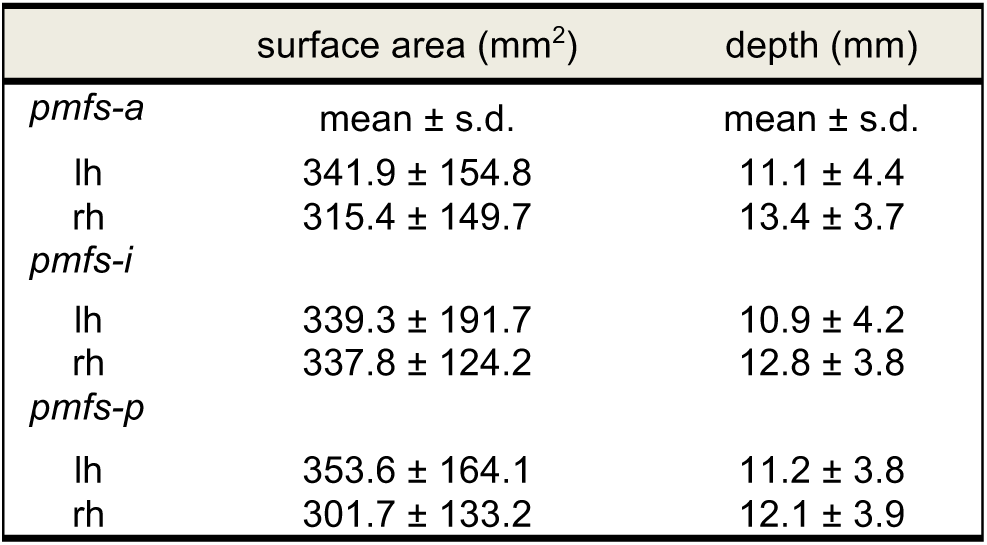
Surface area and depth of the *pmfs*

**Extended Data Figure 5-1.**
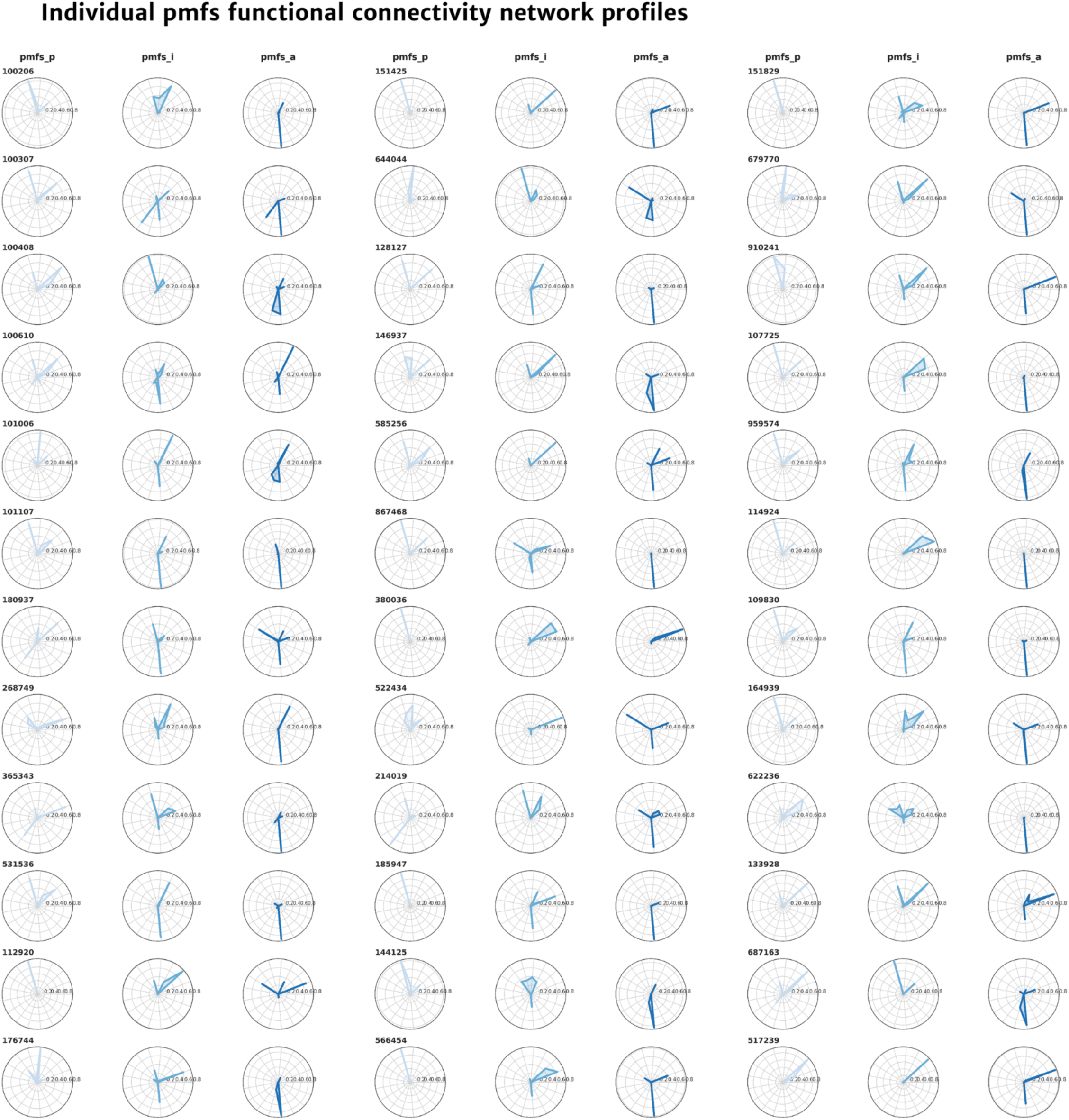
Individual resting-state network connectivity profiles for the *pmfs* components. The individual connectivity profiles and *pmfs* sulcal definitions were used to calculate the connectivity fingerprint, which represents the overlap of each network within the *pmfs* component of each subject. Polar plots showing the connectivity fingerprint of the three *pmfs* components (plotted outwards) with each of 17 resting-state functional connectivity networks for each individual subject (numbered) for the left hemisphere.

**Extended Data Figure 6-1.**
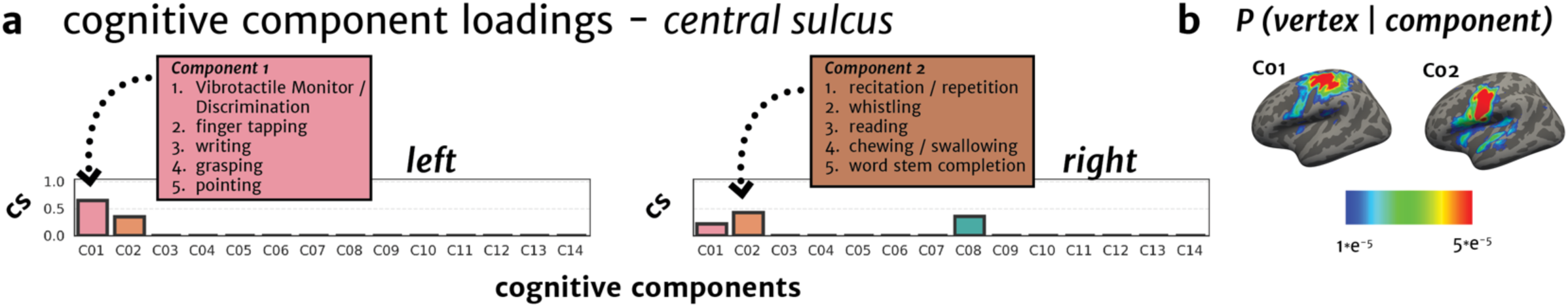
A meta-analysis of fMRI experimental tasks: Cognitive components and the central sulcus (*cs*). (**a**) A posterior probability (y-axis) was calculated for each of 14 cognitive components (x-axis) being associated with the cs (**Materials and Methods**). Somato-motor components of the cognitive component map (C01, C02, right) align most highly with the central sulcus as one would expect, which shows the ability of this method to measure structural-functional correspondences at the meta-analytic level. Descriptions of tasks loading onto C01 and C02 are included in the inset for the left and right hemispheres, respectively. (**b**) Maps of C01 and C02 on the cortical surface showing high overlap with the *cs*.

**Extended Data Figure 7-1.**
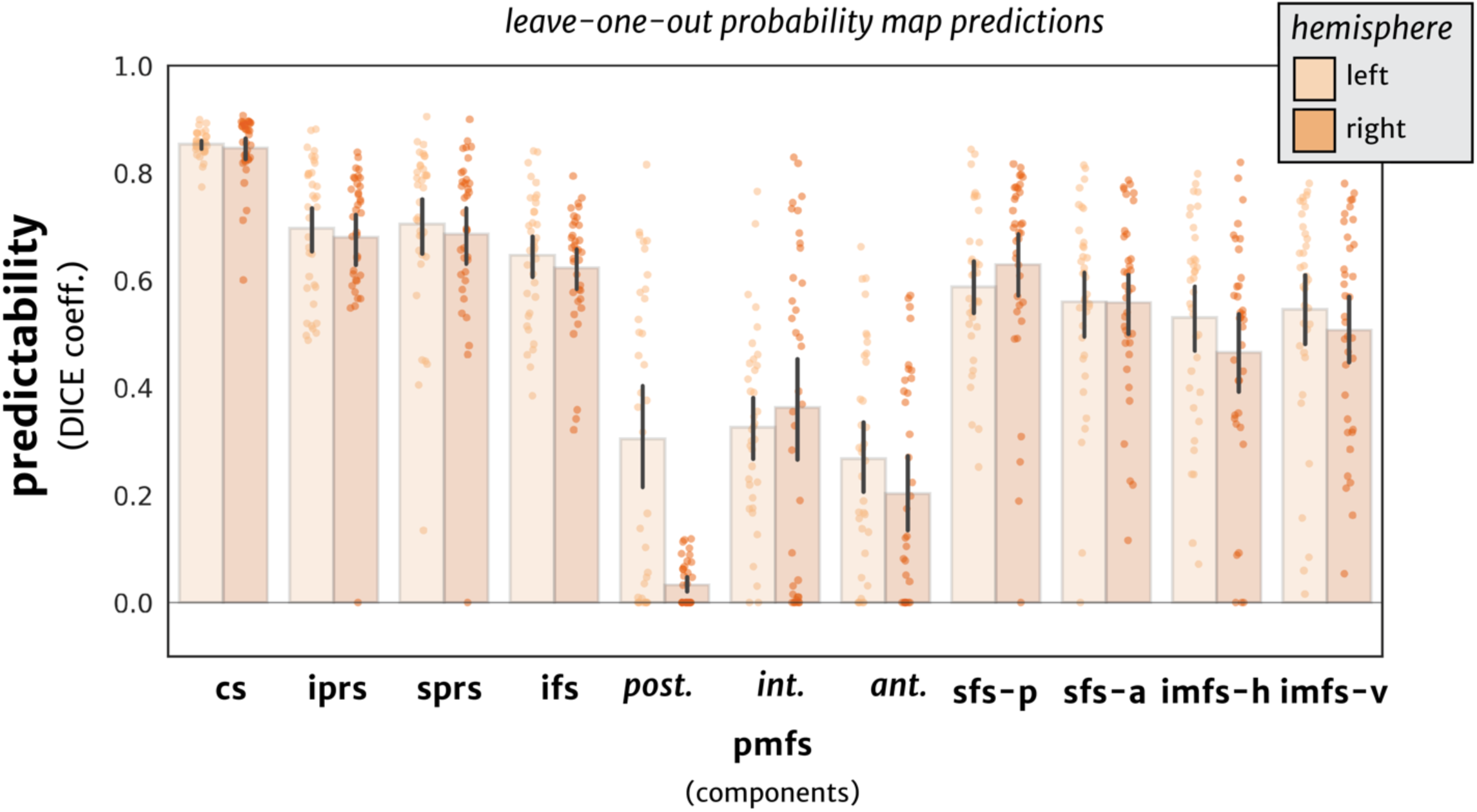
Quantification and predictability for all LPFC sulci. Overlap (DICE coefficient, y-axis) between predicted and manual location of all frontal sulci (x-axis) within individual subjects. Predictability performance is lowest for the *pmfs* components compared to the rest of LPFC sulci.

## Footnotes

1 *To* *Classic neuroanatomy atlases provide schematics of sulci to guide the reader, as well as guide the practice of manual sulcal definitions*. *Modern atlases, including the recent proposal of tertiary sulci that we used to guide the manual definition of LPFC tertiary sulci (Petrides, 2019), continue this historical trend*. *Thus, defining tertiary sulci requires the expertise to apply a two-dimensional schematic of a post-mortem brain to an inflated cortical surface reconstruction*. *This application requires many translational steps including the fact that the schematic does not fully represent individual subject variability – an issue which has been brought up, but remains unresolved, for over a century*. *For example, in reference to his own schematic of sulci and cortical areas, Smith (1907) writes: “*…*no single example is the exact condition represented in these schemata*…*” (Smith, 1907, pg*. *238)*.

2 *To our knowledge, and to quote Klein and colleagues, the “Mindboggle-101 dataset is still the largest publicly available set of manually edited human brain labels in the world” (Klein et al*., *2017)*. *Consisting of 101 brains (202 hemispheres), the dataset contains only 31 labels in each hemisphere*.

